# TEAD1 condensates are transcriptionally inactive storage sites on the pericentromeric heterochromatin in cancer cells

**DOI:** 10.1101/2025.05.02.651992

**Authors:** Yiran Wang, Jindayi Liang, Kimberly S. Lange, Justin Demmerle, Eleanor Liu, Ethan Black, Britney Jiayu He, Christopher J. Ricketts, Shawn R. Yoshida, Shasha Chong, W. Marston Linehan, Jennifer M. Kavran, Chongzhi Zang, Danfeng Cai

## Abstract

TEA domain transcription factor 1 (TEAD1), a Hippo pathway transcription factor important in cellular homeostasis and development, is increasingly implicated in cancer biology. Here, we reveal a novel role for TEAD1 in organizing nuclear condensates, independent of active transcription. Using high-resolution imaging, ChIP-seq, RNA-seq and proximity-based proteomics, we demonstrate that in patient-derived renal cell carcinoma cells, TEAD1 forms micron-sized condensates by binding to the heterochromatic pericentromeric regions using its DNA-binding domain. TEAD-specific MCAT motifs selectively enrich and cluster in the pericentromeric region, specifically seeding TEAD1 condensates. TEAD1 condensates do not activate transcription but instead serve as depots for excess TEAD1, and disrupting TEAD1 condensates leads to increases in YAP/TEAD target gene expression. This organization of TEAD1 contrasts with that observed in other genomic regions of both RCC and normal kidney cells, in which TEAD1 associates with markers of active transcription. Our findings provide a mechanistic framework for TEAD1’s dual regulatory roles, offering new insights into its contribution to transcriptional dysregulation and tumor progression.

## Introduction

Key nuclear processes such as transcription, RNA splicing, and DNA repair occur in membrane-less compartments known as biomolecular condensates^1–3^. These condensates are broadly defined as dynamic assemblies formed through weak interactions between proteins and nucleic acids^1,2,4^. Transcriptional condensates, specifically, are formed by transcription-related factors^5^. Although they have garnered increased attention, their mechanisms of formation and function remain poorly understood. While liquid–liquid phase separation is the dominant model explaining their assembly^6,7^, alternative mechanisms—such as low-valency interactions with spatially clustered DNA binding sites—have also been proposed^8,9^. Moreover, although transcriptional condensates are generally associated with promoting gene expression^10–12^, some have been shown to repress transcription^13,14^. Thus, a deeper understanding of how transcriptional condensates form and function in biologically relevant contexts is essential, particularly for developing strategies to target them in diseases like neurodegeneration and cancer^15,16^.

Renal cell carcinoma (RCC) is a common type of kidney cancer. While there have been considerable improvements in disease-free survival among patients with RCC, effectively managing metastatic disease continues to be a significant challenge in clinical practice, and a clear understanding of the molecular mechanism of RCC progression is still much needed^17^. Genetic alterations in RCC frequently occur in the Hippo pathway^18^, a deeply conserved signaling network regulating cell growth^19^. The Hippo pathway includes key proteins such as STE20-like kinases 1/2 (MST1/2), large tumor suppressor kinases 1/2 (LATS1/2), scaffold proteins such as Salvador homologue 1 (SAV1) and neurofibromatosis type 2 (NF2), which can be activated by cell-cell contacts^20^. The main role of the Hippo pathway is to phosphorylate and repress its downstream transcriptional coactivators, Yes-associated protein (YAP) or its paralog, transcriptional coactivator with PDZ-binding motif (TAZ), by sequestration in the cytoplasm^21–23^.

Therefore, when the Hippo pathway is “off” or mutated such as in RCC (most mutations occur in NF2 or SAV1^18,24^), YAP/TAZ translocate into the nucleus, bind to transcription factor TEA domain 1-4 (TEAD1-4), and activate genes promoting cell proliferation and survival^25^. However, where and how the Hippo pathway components are activated inside the cell is not well understood.

Recently, many components of the Hippo pathway have been found to form biomolecular condensates. The two transcription coactivators, YAP and TAZ, were first to be discovered to form condensates that activate transcription^4,10,26^. Subsequently, upstream Hippo pathway kinases and adaptor proteins were also observed to form condensates and the effects of these condensates on signal transduction varied^27,28^. While transcription factors TEAD1-4 are mostly considered client proteins that are incorporated into existing YAP and TAZ condensates^4,10,26^, a recent study reported that TEAD condensates could turn into active hubs upon YAP/TAZ overexpression^29^. However, the overexpression system used to activate transcription limited understanding the physiological roles of TEAD condensates. Additionally, the function of condensates comprised of Hippo pathway components in various cancers is also poorly understood.

In this study, we focus on transcription condensates formed by TEA domain family member 1 (TEAD1) in Hippo pathway-mutated RCC. We found that, different from most other transcriptional condensates, TEAD1 condensates localize to heterochromatin in the pericentromeric regions and do not mediate active transcription. Our combined ChIP-seq/RNA-seq analysis and proteomics data suggest that TEAD1 condensates may be storage sites that buffer excessive TEAD1, such as in Hippo-mutated kidney cancer cells. These results reveal an interesting alternative function for transcriptional condensates and new ways cells can harness condensates in homeostasis and diseases.

## Results

### TEAD1 forms large condensates in cancer cells

We aimed to determine if endogenous YAP or TEAD forms biomolecular condensates in cancer cells, especially in those with high YAP or TEAD expression. Therefore, we examined a previously established diverse panel of patient-derived RCC cells, the UOK cell lines^30^. Among these lines, UOK121, UOK122, UOK139, and UOK140 are clear cell RCC lines, and UOK275 and UOK342 are papillary RCC lines. A non-cancerous kidney tubule cell line, HK2, was used as a control. Since phase separation and condensate formation are tightly coupled to protein expression levels^1^, we first checked the levels of YAP and TEAD1 in these cell lines. We found that HK2, UOK121, UOK342, UOK122, and UOK140 expressed high levels of YAP (**Fig. 1a,b**). Among these cell lines, UOK121 and UOK342 also expressed high levels of TEAD1 (**Fig. 1a,b**). Examining these cell lines for YAP/TEAD1 condensate formation, we found that while YAP is diffusely localized to the nucleus and forms small condensates in different cell lines (**Extended Data Fig. 1h**), TEAD1 forms prominent micron-sized condensates only in TEAD1-high UOK121 and UOK342 cells (**Fig. 1c**). We observed at least one, and up to five, TEAD1 condensates per nucleus in the majority of both UOK121 and UOK342 cells (**Fig. 1d**); most of these TEAD1 condensates were either close to the nuclear lamina or the nucleolus (**Fig. 1e,f**). These condensates were neither artifacts of immunofluorescence (IF), as TEAD1 knockdown by RNA interference (RNAi, **Extended Data Fig. 1a,b**) eliminated TEAD1 condensates (**Extended Data Fig. 1c**), nor aggregates as they did not stain positively for an amyloid marker AmyTracker (**Extended Data Fig. 1d**)^31^. Interestingly, TEAD1 condensates appear to be a general feature of cancer cells; a panel of melanoma, breast cancer, cervical cancer, and osteosarcoma cell lines mostly contained TEAD1 condensates while the respective noncancerous cells did not (**Extended Data Fig. 2a-e**). Among other members of the TEAD family, TEAD2 and TEAD3 do not form condensates, while TEAD4 does form condensates, although less prominently, in UOK121 and UOK342 cell lines (**Extended Data Fig. 1e**). These results may be explained by the lower expression of TEAD2 and TEAD3 than TEAD1 and TEAD4 in different cell lines (**Extended Data Fig. 1f,g**). We have previously found that TEAD1 cannot form condensates by itself but can incorporate into YAP condensates and stabilize YAP condensate formation^10,31^. Interestingly, we here find that TEAD1 forms condensates that do not enrich endogenous YAP (**Extended Data Fig. 1h,i**). We were intrigued by these TEAD1 condensates and went on to examine how they form and function.

**Figure 1.**
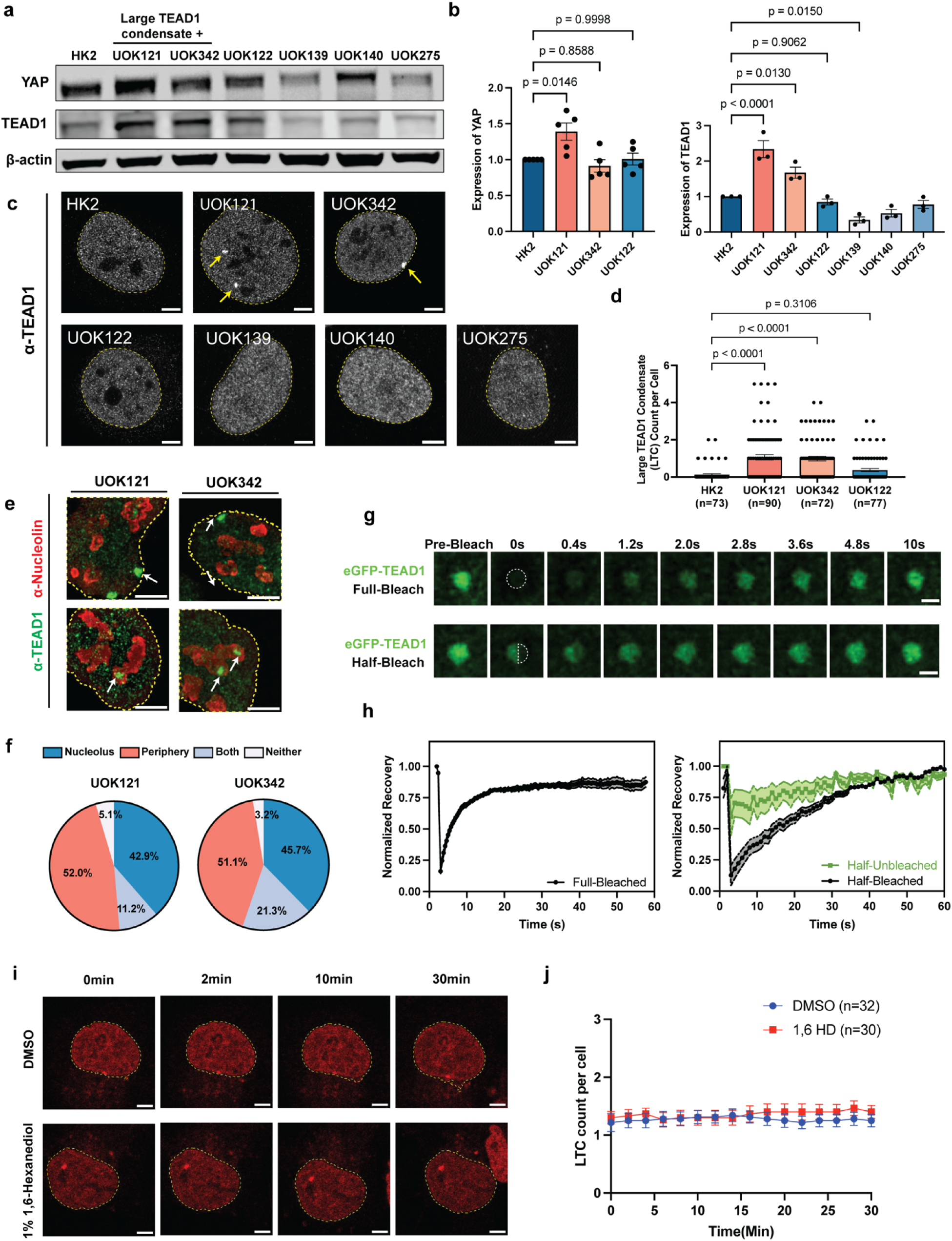
TEAD1 forms condensates in various RCC lines. **(a)** Representative immunoblot showing TEAD1 and YAP protein levels in large TEAD1 condensate (LTC)-positive versus LTC-negative cell lines. **(b)** Quantification of TEAD1 and YAP expression bands from **(a)**, normalized to loading control. Data are presented as mean ± SEM. **(c)** Immunofluorescence imaging demonstrating TEAD1 condensates formation in kidney tubule and renal carcinoma cell lines. Arrows indicate the TEAD1 condensates. Scale bar = 5 μm. Nuclear boundaries are indicated by dotted lines. **(d)** Quantification of large TEAD1 condensate (LTC) in UOK121, UOK122, UOK342, and HK2 cell lines using 3D analysis. Data are presented as mean ± SEM. **(e-f)** Maximum intensity projection from 3D images of TEAD1 (green) and Nucleolin (red) co-immunofluorescence in UOK342 and UOK121 **(e),** with quantification of TEAD1 condensates nucleolar and perinuclear localization (n= 94-98 cells) **(f)**. **(g)** Full-bleach (top) Fluorescence Recovery after Photobleaching (FRAP) and half-bleach FRAP (bottom) of EGFP-TEAD1 UOK342 stable cell line over 10s. Scale bar = 2 μm **(h)** Quantification of recovered fluorescence intensity of fully bleached TEAD1 condensates (left, blue), half-bleached region (right, blue), and unbleached region (right, magenta). Shaded areas indicate the standard error of the mean (SEM) for each measurement (n = 10). **(i-j)** Representative live-cell image of UOK342 TEAD1–HaloTag knock-in cells after treatment with DMSO control (top) or 1,6-hexanediol (1,6HD, bottom). HaloTag fluorescence was labeled with JFX554 HaloTag-ligand. Quantification of LTC count per cell shown in **(j)**. Scale bar = 5 μm. Data are shown as mean ± SEM. Statistical significance of all data was determined by one-way ANOVA, only p-values <0.05 are shown.

To probe the biophysical properties of TEAD1 condensates, we employed fluorescence recovery after photobleaching (FRAP). Specifically, we used full-condensate FRAP to examine the diffusion of TEAD1 molecules between TEAD1 condensates and the nucleoplasm, and half-condensates FRAP to examine the dynamics of TEAD1 inside the condensates. We found that TEAD1 condensates recovered fluorescence rapidly after full-condensate FRAP (t_1/2_ = 3.44 ± 0.067 seconds, mobile fraction = 0.838), indicating that TEAD1 exchanges quickly between the nucleoplasm and the condensates (**Fig. 1g, 1h left**). In half-condensate FRAP experiments, the bleached area also recovered fluorescence rapidly (**Fig. 1g, 1h right, black line**), accompanied by a 25% dip in the unbleached area immediately after bleaching (**Fig. 1h right, green line**). These findings indicate that TEAD1 molecules preferentially mix within TEAD1 condensates, consistent with an LLPS-driven mechanism^8^. Treatment with 1,6-hexanediol (1,6HD), an aliphatic alcohol^1,2,32^, on the other hand, did not dissolve the TEAD1 condensates in the nucleus nor did it alter their number (**Fig. 1i,j**). Together, these results all indicate that TEAD1 condensates are liquid-like compartments stabilized by interactions that are not primarily hydrophobic.

### TEAD1 condensates do not mediate active transcription

TEAD family transcription factors usually mediate active gene expression. To investigate if TEAD1 condensates also mediate active transcription, we examined their association with either RNA polymerase II phosphor-Serine2 (RNA Pol II-pSer2), a marker of transcription elongation^33^, H3K27 acetylation (H3K27ac) — a histone modification denoting active enhancers^34^, or Bromodomain-containing protein 4 (BRD4) — a transcriptional coactivator found at super-enhancers^35,36^. To our surprise, TEAD1 condensates did not colocalize with any of these active transcription markers but instead excluded them (**Fig. 2a-b**). Consistently, there was no active transcription from large TEAD1 condensates as evidenced by a lack of 5-ethynyluridine (EU) staining in nascent RNA labeling experiments, unlike what was seen for the nucleolus, an active ribosomal RNA transcription center (**Fig. 2c,d, Extended Data Fig. 3a**). Furthermore, TEAD1 condensates did not colocalize with the canonical YAP/TEAD target gene *CYR61* (also called *CCN1*)^37,38^ when visualized by simultaneous IF against TEAD1 and nascent RNA-FISH against *CYR61* (**Fig. 2e-g**). These results indicate that TEAD1 condensates do not mediate active transcription at the YAP/TEAD1 canonical target gene. In addition, TEAD1 condensates did not colocalize with other known nuclear membrane-less organelles (MLOs) such as Cajal bodies, nuclear speckles, or PML nuclear bodies (**Extended Data Fig. 3b-e**). TEAD1 condensates therefore represent a unique kind of condensate that contains a transcription factor yet does not mediate active transcription. Interestingly, we found that TEAD1 not only forms large TEAD1 condensates (LTC) but also forms many small TEAD1 condensates (STC). Compared to LTC (1.31 +-0.48 µm^3^ in size and around 3 per cell), the STC are only 0.030+-0.015 µm^3^ in size and average ∼169 per cell (**Fig. 2h**). The STC resemble canonical transcriptional condensates: they colocalize with coactivator YAP and mediate active transcription evidenced by their colocalization with YAP/TEAD1 target gene transcript *CYR61* (**Fig. 2i,j**), and their much shorter distance to *CYR61* than the LTC (**Fig. 2k,l**). Therefore, TEAD1 can form two types of condensates that differ in size and functions. We will focus on characterizing the large TEAD1 condensates as they differ from canonical transcription condensates and refer to them below as TEAD1 condensates for simplicity.

**Figure 2.**
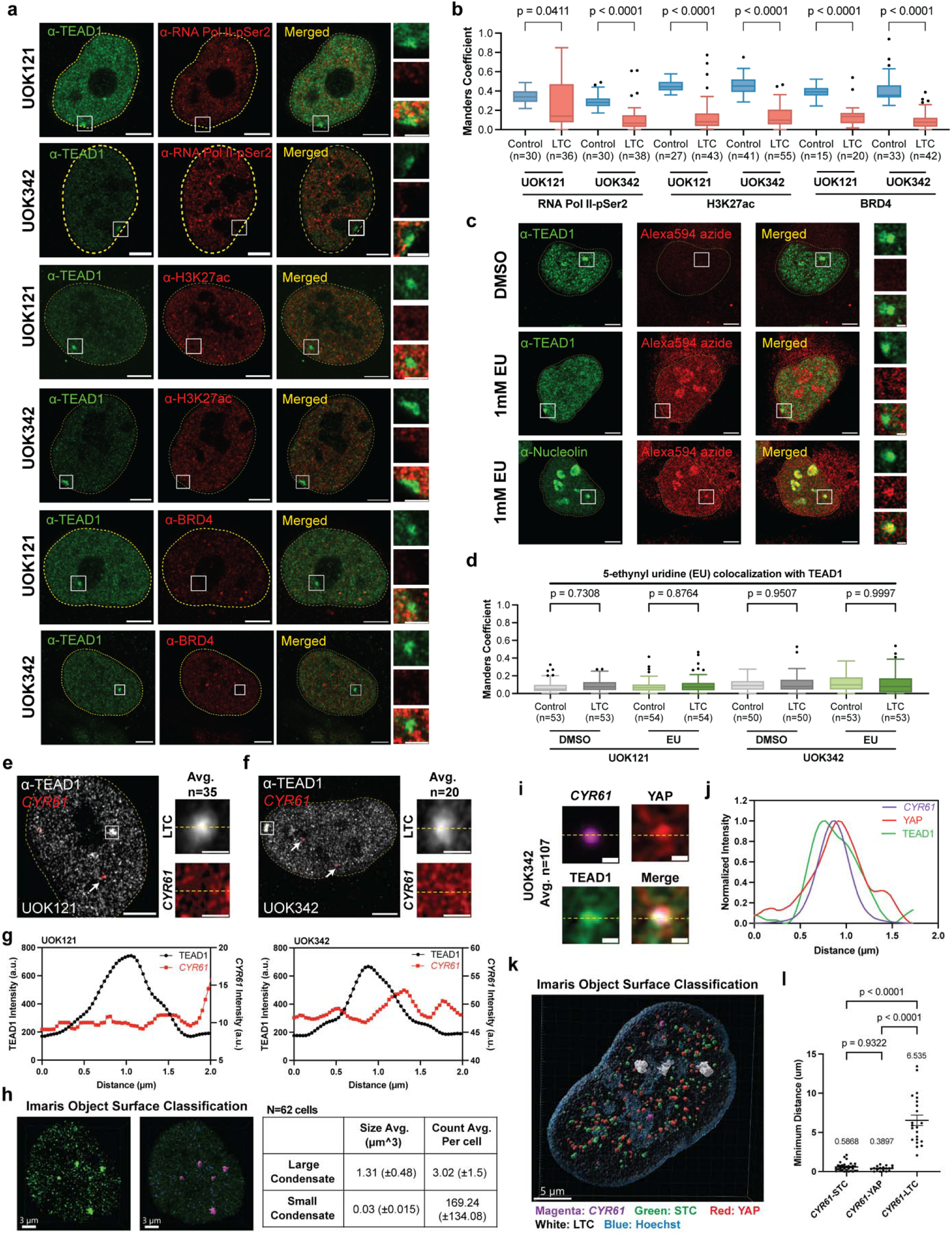
Large TEAD1 condensates are transcriptionally inactive. **(a)** Representative immunofluorescence images of TEAD condensates (green) and active transcription markers (red) RNAPII2, H3K27ac, and BRD4 in UOK121 and UOK342. Scale bars = 5µm. Insets show magnified boxed regions of merged channels with adjusted intensity. Scale bars = 2 μm. Nuclear boundaries indicated by dashed lines. **(b)** Quantitative assessment and statistical evaluation of co-localization of TEAD1 with active transcription markers around condensates and control regions. Data are presented as mean ± SEM. **(c)** Representative immunofluorescence images of TEAD1 (top) and Nucleolin (bottom) in UOK342 treated with DMSO control or 1mM 5-Ethynyl Uridine (EU) for 10min. Nascent RNA incorporated with EU are visualized by Click-iT™ RNA Alexa Fluor 594 azide. Scale bars = 5µm **(d)** Quantification of colocalization of TEAD1 with EU in UOK121 and UOK342 treated with DMSO control or EU for 10min, measurement made around LTC and control regions. **(e-f)** Representative images of large TEAD1 condensate (LTC) and Fluorescence in Situ Hybridization (FISH) of nascent *CYR61* mRNA in UOK121 **(e)** and UOK342 **(f)**. Arrows indicating *CYR61* nascent mRNA signals and TEAD1 condensates are highlighted in boxed regions. Nuclear boundaries are indicated by dotted lines. Insets show averaged images centered on large TEAD1 condensate (LTC), TEAD1 (gray) and *CYR61* (red), in UOK121 (top, n=35) and UOK342 (bottom, n=20). Scale bars = 1µm. **(e)** Line plots of large TEAD1 condensates (LTC) and *CYR61* average intensity along the dashed line shown for UOK121 (left) and UOK342 (right). **(f)** Representative images of small TEAD1 condensate (Blue) and large TEAD1 condensate (LTC, pink) 3D surface identification on Imaris. Table on the right shows average size and count of two types of condensates in UOK121 and UOK342. N=62 cells. **(g)** Averaged images centered on nascent *CYR61* mRNA, *CYR61* (magenta), YAP (red), and TEAD1 (green) in UOK342 (n=107). Scale bars = 0.5µm. **(h)** Line plots of *CYR61*, YAP and TEAD1 average intensity along the dashed line shown on **(i)**. **(i)** Representative images of 3D Object Surface Classification processed on Imaris 10.2.0 (Bitplane). **(j)** Quantification of minimum distance from nascent *CYR61* mRNA to STC, YAP, and LTC. N=9-17 cells. Statistical significance determined by one-way ANOVA.

### TEAD1 condensates localize to the pericentromeric heterochromatin region

Since TEAD1 condensates do not associate with markers of active transcription (**Fig. 2a-b**) and are close to the nuclear lamina and the nucleolus (**Fig. 1e-f**), regions typically enriched in heterochromatin, we went on to test if TEAD1 condensates localize to the heterochromatin. Histone 3 lysine 9 trimethylation (H3K9me3) is an epigenetic marker characteristic of heterochromatin^39^. Co-staining of TEAD1 and H3K9me3 showed that TEAD1 condensate colocalizes with an H3K9me3 domain in the TEAD1 condensates-positive kidney cancer and other cancer cell lines (**Fig. 3a-d, Extended Data Fig. 2f**). No co-occupancy between TEAD1 and H3K9me3 was observed in the TEAD1 condensates-negative UOK122 or HK2 cells (**Extended Data Fig. 4a-d**). In addition, in UOK121 cells, TEAD1 condensates also colocalize with heterochromatin protein 1 alpha (HP1α, **Extended Data Fig. 4e-h**), a protein responsible for heterochromatin assembly^40^. Polycomb bodies organize heterochromatin regions to repress gene expression^41,42^. TEAD1 condensate did not colocalize with markers of Polycomb bodies (**Extended Data Fig. 4i, j**), indicating that TEAD1 condensates localize at unique heterochromatin regions distinct from Polycomb bodies. Instead, TEAD1 condensates were always adjacent to the centromere, showing bimodal distribution around the centromere together with H3K9me3 (**Fig. 3e-h, Extended Data Fig. 4k-n**), indicating that TEAD1 condensates specifically bind to the pericentromeric heterochromatin, a region crucial for centromere integrity and proper segregation of chromosomes during mitosis^43,44^. This is further supported by the much closer distance of TEAD1 condensates to the centromere histone marker centromere-specific protein A (CENP-A, 0.2 µm)^43^ than to the telomere marker telomeric repeat binding factor 2 (TRF-2, 1 µm, **Fig. 3i-l**)^45^. These results indicate that TEAD1 condensates specifically localize to the pericentromeric heterochromatin.

**Figure 3.**
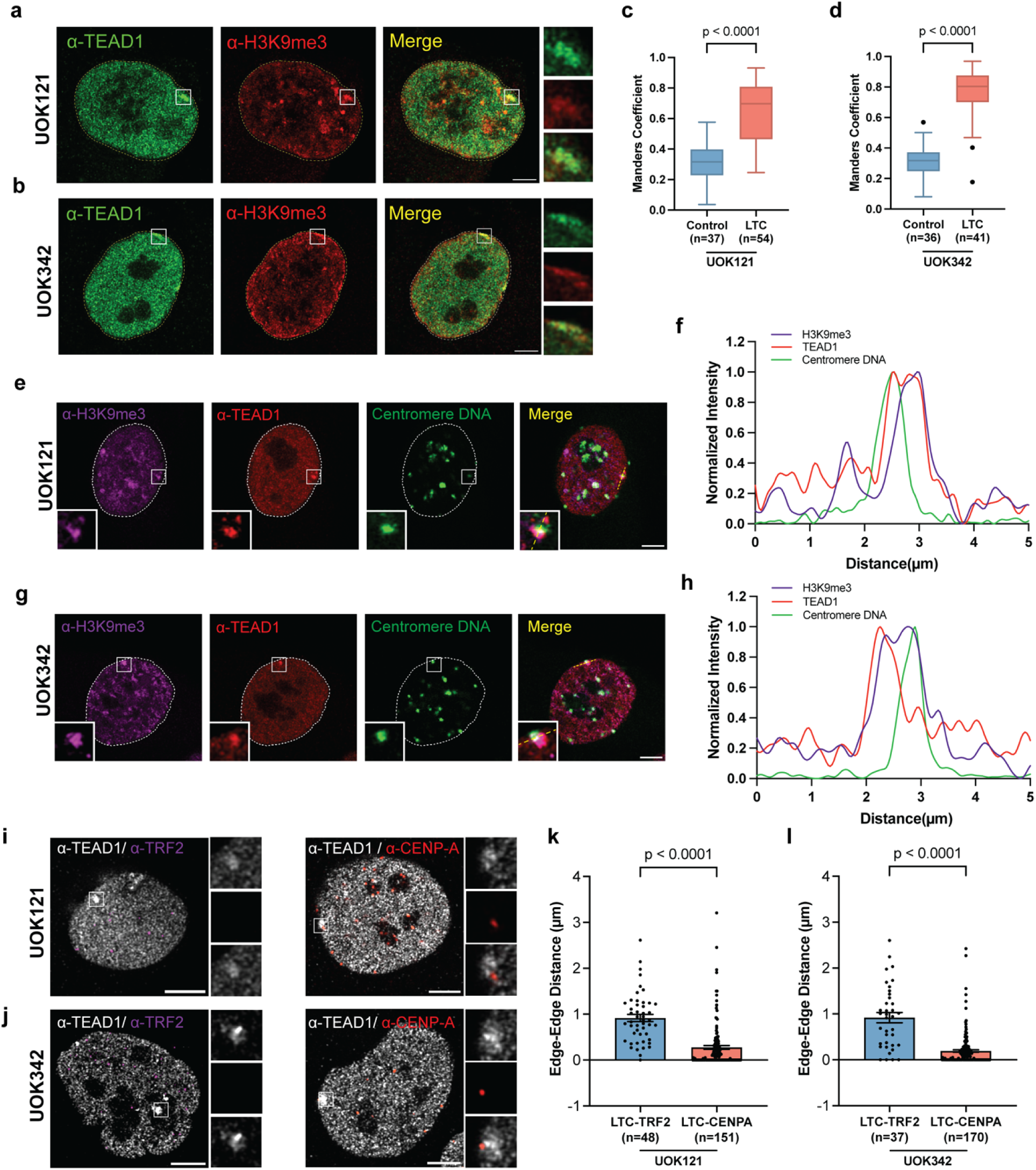
TEAD1 condensates colocalize with heterochromatin markers in different RCC cell lines. **(a-d)** Representative immunofluorescence images of TEAD1 (green) and H3K9me3 (red) in UOK121 **(a)** and UOK342 **(b)** cells (scale bars are 5μm). Insets show magnified images of boxed regions. Co-localization of TEAD1 condensates and H3K9me3 in UOK121**(c)** and UOK342**(d)** compared to the more diffused TEAD1 region. **(e-h)** Representative immunofluorescence images illustrate the co-localization of TEAD1 (red) with H3K9me3 (magenta) and adjacent to centromere DNA-FISH (green) and in UOK121 **(e)** and UOK342 **(g)** cells. Insets show magnified images of boxed regions. Scale bars are 5µm. Signal intensity profiles along the lines indicated on the merged-channel images for UOK121 **(f)** and UOK342 **(h)**. **(i-l)** Representative immunofluorescence images and quantification illustrate the co-localization of TEAD1 (green) with TRF2 (magenta) and CENP-A markers (red) in UOK121 **(i)** and UOK342 **(j)**. Insets show magnified images of boxed regions. Quantification uses 3D nearest neighbor analysis in UOK121 **(k)** and UOK342 **(l)**, demonstrating a tighter distribution of CENP-A at a smaller minimal distance. Data are presented as mean ± SEM. Statistical significance determined by unpaired t-test.

### DNA binding domain of TEAD1 is essential for TEAD1 condensate formation

The localization of large TEAD1 condensates to the pericentromeric region suggests that TEAD1 condensates formation may be seeded by DNA binding. To test this possibility, we generated variants of TEAD1 with impaired DNA binding. The DNA-binding domain of TEAD1 (TEA domain) is comprised of a highly conserved homeodomain region containing three α-helices, Helix 1 (H1), Helix 2 (H2), and Helix 3 (H3)^46^ (**Fig. 4a**). To disrupt DNA binding by TEAD1, we constructed TEAD1 variants that either lacked the DNA binding domain (Δ55-122) or bore previously validated site-specific substitutions in the DNA binding domain (either F43P and L47P, TEAD1H1Mut or Q89P and S92P, TEAD1H3Mut) (**Fig. 4a**)^47^. When transiently expressed in both LTC-positive and negative cells, we found that compared with EGFP-TEAD1 WT, variants with impaired DNA binding formed fewer condensates (**Fig. 4b-e, Extended Data Fig. 5a, b**) under similar or even higher expression levels (**Extended Data Fig. 5c-g**). In addition, the GFP signal of these mutants did not colocalize with the heterochromatin (**Fig. 4f-k, Extended Data Fig. 5h-m**). On the other hand, altering other TEAD1 regions such as the N-terminal intrinsically disordered region (IDR), YAP-binding site (Y406)^48^, or the protein kinase A phosphorylation site (S102)^49^ did not disrupt large TEAD1 condensates nor their association with the heterochromatin (**Fig. 4a-e, l-p**). These data indicate that the DNA-binding domain of TEAD1 is essential for TEAD1 condensates formation on the heterochromatin.

**Figure 4.**
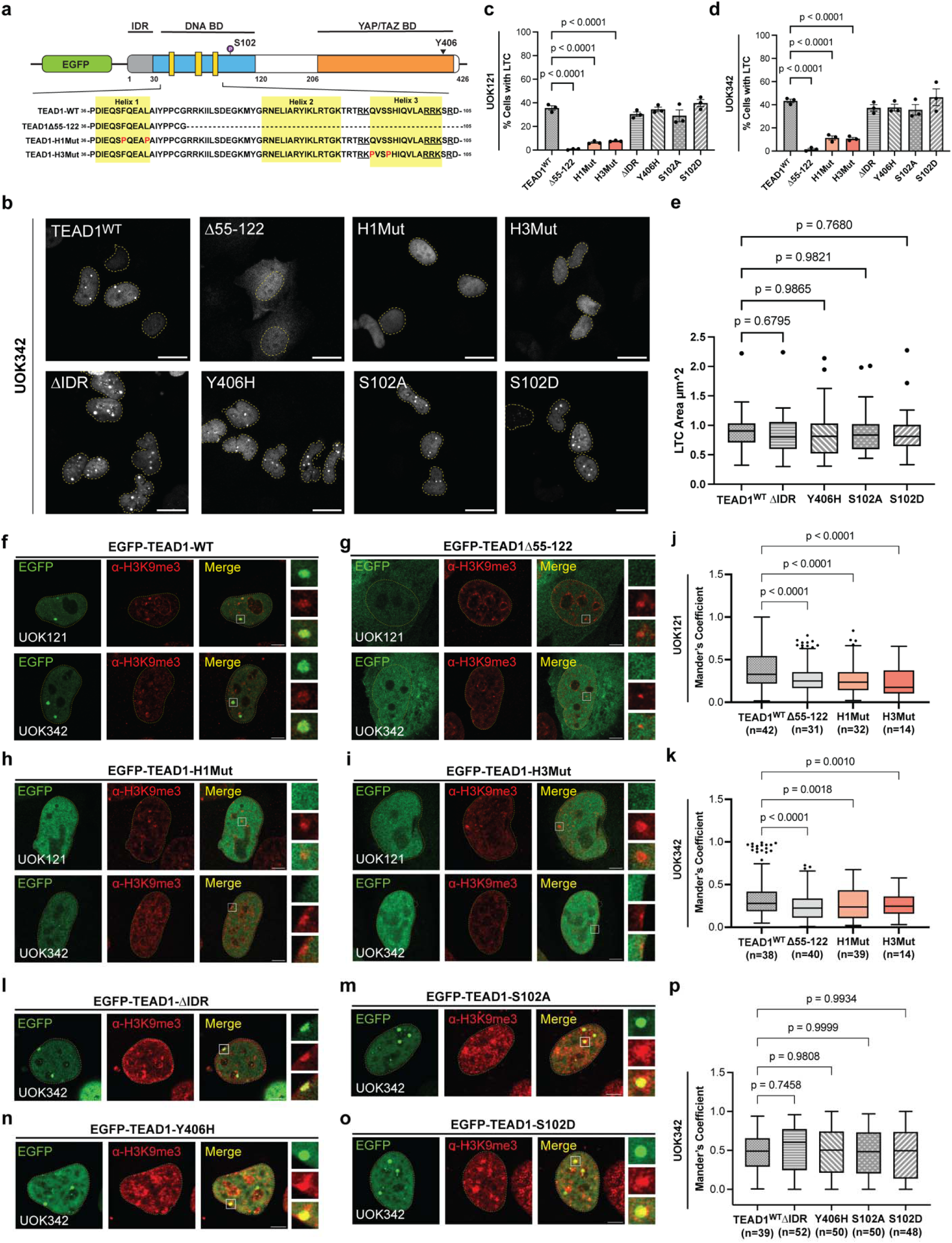
TEAD1 condensates formation is mediated by TEAD1 DNA-binding domain. **(a)** Illustration of expression constructs of EGFP-tagged wild-type TEAD1 and DNA-binding domain (DNA BD) mutants. The underlined amino acid sequence represented the TEAD1 nuclear localization signal (NLS), and the red amino acids are mutated in the respective mutants. PKA phosphorylation site labeled in purple(S102). Triangle dot indicates site responsible for YAP binding(Y406). **(b-d)** Representative images of UOK342 cells expressing EGFP-tagged TEAD1 wild-type and mutants in **(b)**. Scale bar = 20µm. Quantification of percentage LTC-positive cells upon transfection in UOK121 **(c)**, and UOK342 **(d)**. Data are presented as mean ± SEM. **(e)** Quantification of LTC area of UOK342 cells expressing EGFP-TEAD1-WT and LTC-positive mutants shown in **(b-d)**. **(f-i)** Representative images of UOK121 and UOK342 cells expressing EGFP-TEAD1-WT **(f)**, EGFP-TEAD1-DNA BD deletion **(g)**, EGFP-TEAD1-H1Mut **(h)**, and EGFP-TEAD1-H3Mut **(i)**, with immunofluorescence signal of H3K9me3 (red). Insets show magnified images of boxed regions. Scale bars = 5μm. **(j-k)** Quantification of co-localization between H3K9me3 and EGFP-TEAD1 variants shown in **(f-i)**. **(l-o)** Representative images of UOK342 cells expressing EGFP-TEAD1-ΔIDR **(n)**, EGFP-TEAD1-Y406H **(o)**, EGFP-TEAD1-S102A **(p)**, and EGFP-TEAD1-S102D **(q)**, with immunofluorescence signal of H3K9me3 (red). Insets show magnified images of boxed regions. Scale bars = 5μm. **(p)** Quantification of co-localization between H3K9me3 and EGFP-TEAD1 variants shown in **(l-o)**. Statistical significance determined by one-way ANOVA.

### TEAD1 and YAP co-occupy distal enhancer sites to upregulate gene expression

To understand the functions of TEAD1 in regions both outside and inside of TEAD1 condensates, we set out to determine the binding sites of TEAD1 on the genome and investigate their relations to gene expression. To achieve this, we performed chromatin immunoprecipitation followed by sequencing (ChIP-seq) using antibodies against endogenous TEAD1, YAP, H3K27ac, as well as RNA sequencing (RNA-seq), in both TEAD1 condensates-positive (UOK121, UOK342) and condensates-negative (UOK122, HK2) cell lines. TEAD1 and YAP are known to co-occupy distal enhancer sites to upregulate gene expression in various stem cells and cancer cells^50–52^. Consistent with these results, we found that in all cell lines, most TEAD1/YAP co-occupied peaks mapped to either distal intergenic (41.9%) or intronic regions (46.7%), while few (5.2%) were found at promoters (**Extended Data Fig. 6a**). Specifically, over 50% of TEAD1/YAP-bound regions were 50kb away from the transcription start sites (TSS) in UOK121 and UOK342 (**Extended Data Fig. 6b, c**). Alignment of these distal TEAD1-bound sites with H3K27ac peaks revealed a bimodal distribution of H3K27ac centered on TEAD1/YAP peaks (**Fig. 5a**). These results indicate that most TEAD1 binds to distal enhancer sites. These regions were also enriched for Gene Ontology (GO) terms such as actin cytoskeleton organization and cell junction assembly (**Extended Data Fig. 6d**), which are typical TEAD1/YAP-regulated pathways. As expected, TEAD1 and YAP peaks colocalize at canonical TEAD1/YAP target genes such as CTGF and CYR61 (**Extended Data Fig. 6e, f**). Combining ChIP-seq and RNA-seq data and using binding and expression target analysis (BETA)^53^, we found that most YAP/TEAD1 co-bound regions have upregulated gene expression (**Extended Data Fig. 6g, h**). Since TEAD1 is a sequence-dependent transcription factor, we performed MEME-ChIP to examine its binding motif. Not surprisingly, TEAD family-specific MCAT motif^54^ is the most enriched binding motifs in regions bound by TEAD1 (**Fig. 5b**).

**Figure 5.**
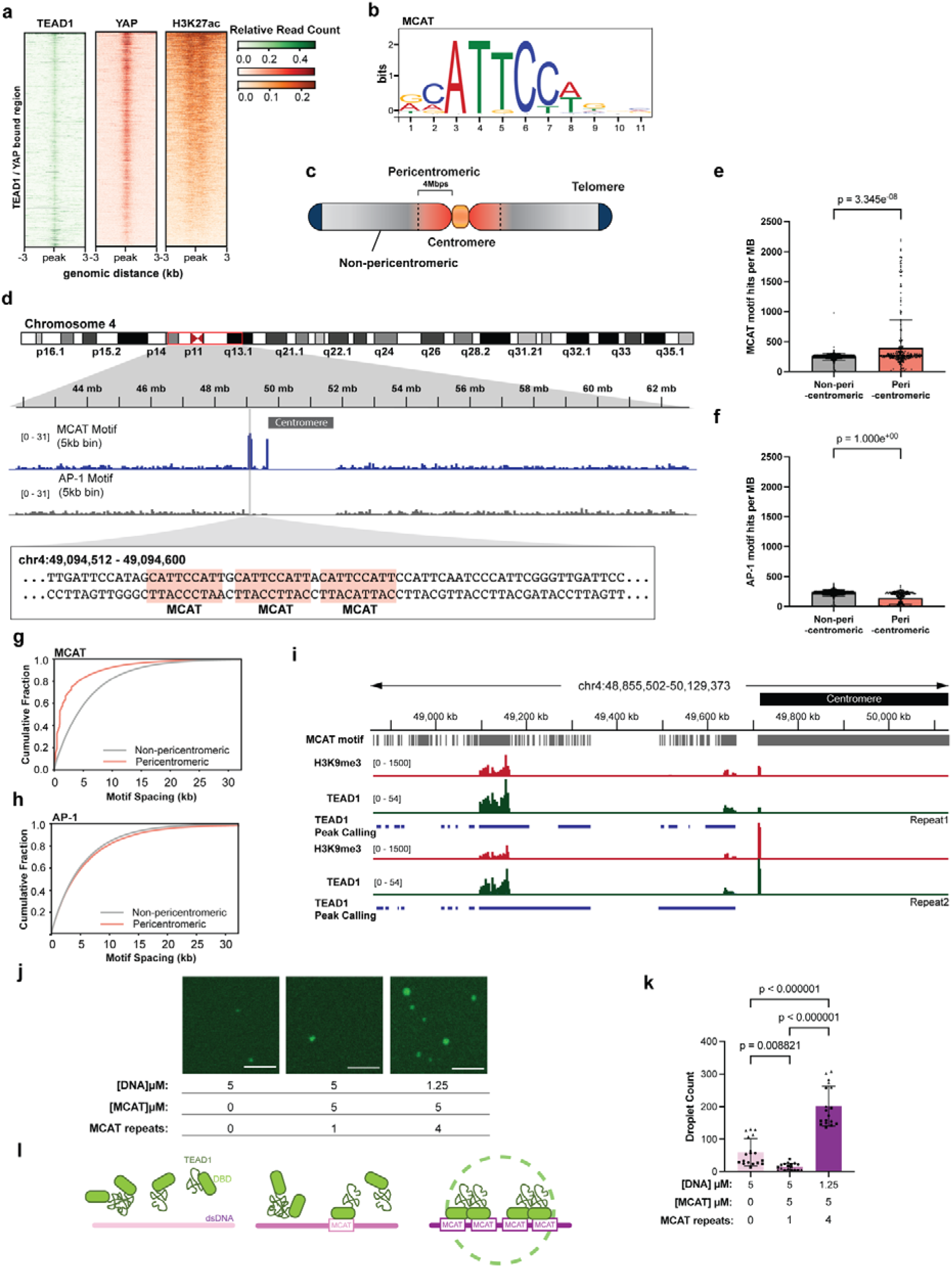
TEAD1 condensates are seeded by the repeated MCAT motifs. **(a)** Heatmaps showing merged TEAD1, and H3K27ac ChIP-seq signals (merged data from HK2). Active enhancer regions are shown as regions that have H3K27ac peaks. **(b)** MEME-ChIP result showing the MCAT motif is present from TEAD1 ChIP data from **(a)**. **(c)** Schematic of chromosome structure. The pericentric region is defined as 4MB flanking the centromere region. **(d)** Gene tracks centered around centromeric and pericentromeric region on chromosome 4 of hg38 reference genome. MCAT and AP-1 motif-count BED files were converted into BigWig density tracks with 5000bp bin size. A small region (chromosome 4:49094512 bp – 49094600 bp) is zoomed in to examine multiple MCAT sequences. Three MCAT motifs are highlighted in red. **(e-f)** Quantification of number of MCAT **(e)** and AP-1 **(f)** motifs per MB within the pericentric and non-pericentric regions. Data are presented as scatterplots with mean ± SEM. Statistical significance determined by Mann–Whitney test. **(g-h)** ECDF plot of kb distance between each MCAT **(g)** and AP-1 **(h)** motifs. Orange line: pericentric region; blue line: non-pericentric region. **(i)** Gene tracks showing multi-mapped TEAD1 (green) and H3K9me3 (red) co-binding at the MCAT-dense (grey) region of near centromere in UOK342. Blue tracks are TEAD1 SICER2 peak-called regions with FDR < 0.05. **(j-l)** 5uM of purified, fluorescently tagged TEAD1 was mixed with the indicated concentrations of dsDNAs with either scrambled sequence (pink), one MCAT repeat (lavender), or four MCAT repeats (purple). **(j)** Representative fluorescent images of TEAD1 condensates. Scale bar corresponds to 5 mm. **(k)** The number of condensates was quantified in each condition. Data was plotted as a superplot of 3 biological replicates, indicated by different shapes, with each data point representing a single image. Significance determined one-way ANOVA. **(i)** Cartoon schematic of TEAD1 phase separation in the presence of different DNA sequences illustrating the factors driving condensates formation in each. TEAD1 is in green. dsDNA is colored as in **(i)**.

### TEAD1 condensates localize to clustered MCAT motifs at the pericentromeric heterochromatin

Having established that TEAD1 binds MCAT motifs to activate canonical YAP/TEAD1 target genes, we next examined where TEAD1 condensates form in the genome. Consistent with our imaging data showing that TEAD1 condensates assemble at pericentromeric heterochromatin in a DNA-binding–dependent manner, we found a striking enrichment of MCAT motifs within pericentromeric regions, where these motifs were also highly clustered (**Fig. 5c-e, g**). In contrast, a length-matched AP-1 motif showed neither enrichment nor clustering in these regions (**Fig. 5d, f, h**), indicating that this pattern is specific to TEAD1-binding sites. These results suggest that clustered MCAT motifs at pericentromeric heterochromatin provide a genomic scaffold for TEAD1 condensate nucleation. Consistent with this model, additional ChIP-seq for H3K9me3 revealed that 25%-35% of total TEAD1 binding sites intersected with the H3K9me3 regions, and these TEAD1/H3K9me3 co-bound regions coincide with MCAT-rich pericentromeric regions (**Fig. 5i**, **Extended Data Fig. 7a-d**), indicating that TEAD1 binding to clustered MCAT motifs within heterochromatin drives the formation of TEAD1 condensates.

To define the mechanism of TEAD1 condensate formation, we investigated whether and how TEAD1 interactions with DNA altered TEAD1 solubility and condensate formation. We monitored condensate formation of full-length, purified TEAD1 (**Extended Data Fig. 7e**) incubated with a panel of dsDNA containing differing numbers of MCAT motifs, either 0, 1, or 4 tandem repeats (0xMCAT, 1xMCAT, 4xMCAT) (**Fig. 5j, k**). While TEAD1 formed condensates in the presence of all the DNAs screened, the number of condensates differed suggesting interactions with DNA influence TEAD1 solubility. TEAD1 formed the most condensates in the presence of 4xMCAT DNA, (e.g. 3.4-fold higher than 0xMCATs and 13.7-fold higher than 1xMCAT). In the presence of a single MCAT motif, TEAD1 formed fewer condensates than either 4xMCAT or 0XMCAT. This altered solubility profile supports a model in which TEAD1 condensation is less likely in conditions that support weak or no DNA binding (e.g. 0xMCAT or 1xMCAT), driven mostly by weak TEAD1 self-associations, and in conditions with high affinity DNA binding (e.g. 4xMCAT or pericentromeric regions) condensation is further enhanced by both multi-valent interactions (both protein:protein and protein:DNA) and an increase effective local concentration of TEAD1 (**Fig. 5l**)^46,55^. These cooperative effects drive phase separation of TEAD1–dsDNA complexes beyond the intrinsic phase separation capacity of TEAD1. The effect of multi-valent DNA interactions provides a biophysical rationale for the formation of large TEAD1 condensates in the pericentromeric regions.

### TEAD1 condensates serve as depots to buffer excess TEAD1

To further evaluate whether TEAD1 condensates are associated with transcriptional regulation, we performed differential expression analysis of RNA-seq data comparing condensate-positive kidney cancer cell lines (UOK121 and UOK342) with the condensate-negative HK2 kidney cell line (**Extended Data Fig. 7f, g**)^56^. We next used BETA to determine whether differentially expressed genes were associated with TEAD1–H3K9me3 co-bound genomic regions. TEAD1/H3K9me3 co-binding sites were over 500 kb away from the nearest TSS (**Extended Data Fig. 7h, i**), and genes proximal to these sites showed no significant enrichment among either up-regulated or down-regulated transcripts (**Extended Data Fig. 7j, k**). Consistent with this lack of transcriptional association, increasing the number of TEAD1 condensates from 0 to 3 did not alter overall nuclear TEAD1 abundance or the expression of canonical TEAD target gene *CYR61* (**Fig. 6a-d**). Together with the observation that large TEAD1 condensates are spatially separated from active transcription markers and transcription sites (**Fig. 2**), these data suggest that peri-centromeric TEAD1 condensates are unlikely to directly drive transcription and may instead function as nuclear depots for TEAD1 in kidney cancer cells.

**Figure 6.**
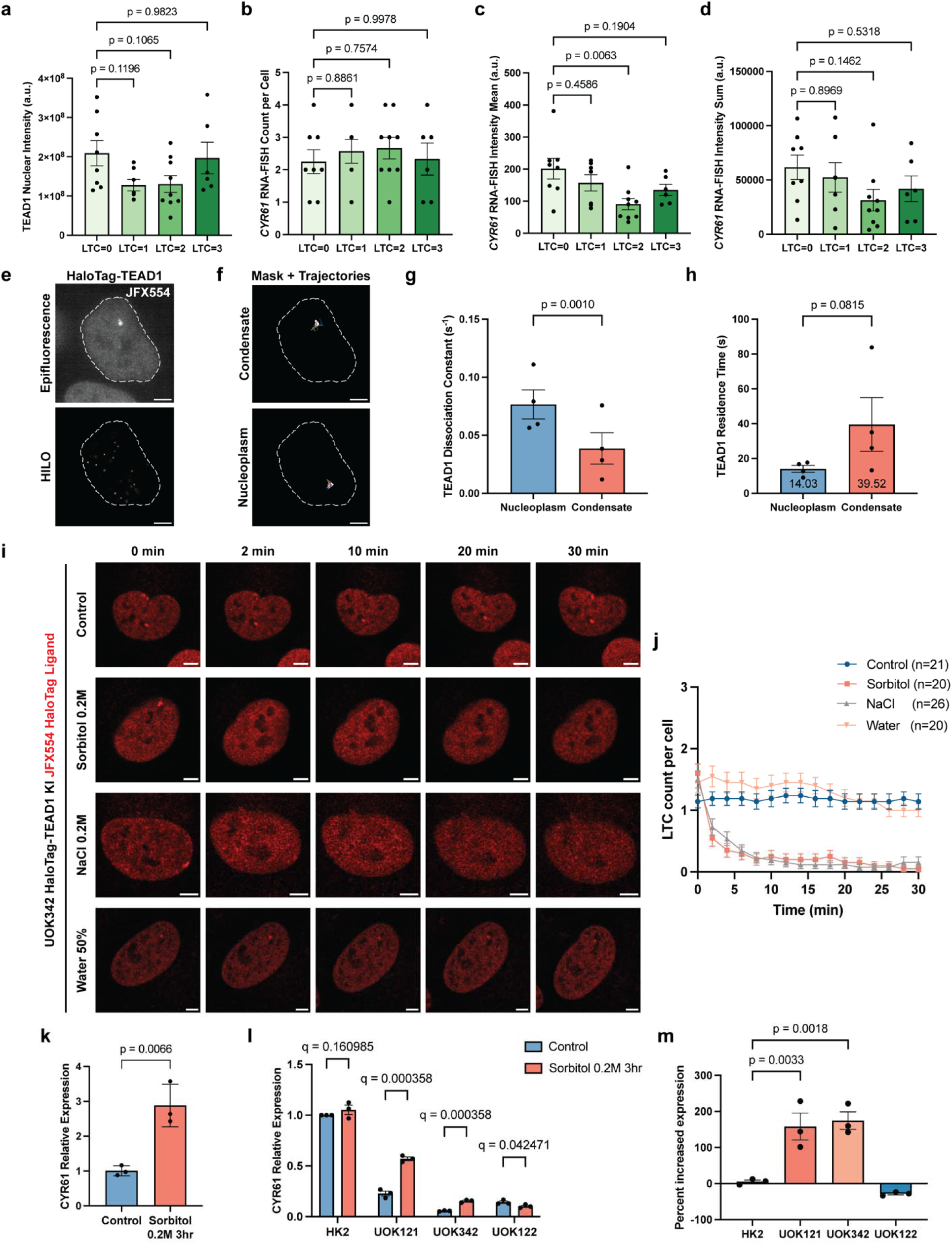
TEAD1 condensates are depots for buffering target gene expression. **(a-d)** Quantification of *CYR61* nascent RNA-FISH with TEAD1 immunofluorescence 3D image analysis. Cell expressing similar level of TEAD1 forms 1-3 large TEAD1 condensates (LTC) in **(a)**. *CYR61* RNA-FISH signal count **(b),** mean intensity **(c)**, and sum intensity **(d)**. **(e)** Representative images of UOK342 HaloTag-TEAD1 KI cell labeled with JFX554 HaloTag ligand for single-particle tracking. Whole nucleus was imaged using epifluorescence microscopy to generate a mask of the TEAD1 condensate (top). HILO microscopy was used for SPT of individual HaloTag-TEAD1 molecules (bottom). Scale bar = 5 μm. **(f)** Trajectories of individual TEAD1 molecules were categorized as inside TEAD1 condensates (top) and or in the nucleoplasm (bottom). To get trajectories located in the nucleoplasm, a binary mask of the TEAD1 condensate generated from **(e)** was randomly rotated and placed elsewhere within the nucleus. Individual trajectories present either in the condensate mask or in the masked area placed randomly in the nucleoplasm are shown on the right, with each color representing a different trajectory. Scale bar = 5 μm. **(g-h)** Quantification of dissociation constants **(g)** and mean residence time **(h)** of individual TEAD1 molecules inside hubs and those outside of hubs in the nucleoplasm. Points represent trajectories grouped by replicate, 8 cells per replicate. Data are presented as mean ± SEM. Statistical significance determined by paired t test, one-tailed. **(i)** Representative live-cell images of UOK342 HaloTag-TEAD1 KI cell line labeled with JFX554 HaloTag ligand and treated with sorbitol 0.2M, NaCl 0.2M, 50% water or control. Scale bar = 5 μm. Quantification of LTC count per cell post-treatment **(j)**. Data is shown as mean ± SEM. **(k)** Quantification of expression of *CYR61* upon 0.2M sorbitol treatment for 3 hours in UOK342 HaloTag-TEAD1 KI cell line compared to control. **(l)** Quantification of expression of *CYR61* upon 0.2M sorbitol treatment for 3 hours in LTC-negative cell lines (HK2, UOK122) and LTC-positive cell lines (UOK121, UOK342). Data are presented as mean ± SEM. Statistical significance determined by multiple unpaired t tests. **(m)** Percent increased *CYR61* expression upon sorbitol treatment relative to control of each cell line in **(l)**. Data are presented as mean ± SEM. Statistical significance determined by one-way ANOVA.

In Hippo mutated cancer cells, depots may buffer TEAD1 overexpression so that downstream gene expressions remain stable. Indeed, single particle tracking experiments showed that TEAD1 dissociated later, and stayed longer within large TEAD1 condensates than in the nucleoplasm (**Fig. 6e-h**), suggesting that large TEAD1 condensates trap TEAD1 proteins within. To definitively show that large TEAD1 condensates serve as depots, we sought for ways to disrupt TEAD1 condensates without mutating the TEAD1 DNA - binding domain and measured its effect on TEAD1 target gene expression. Hyperosmotic stress mediated by sorbitol or NaCl rapidly dissociated large TEAD1 condensates, while hypo-osmotic stress did not (**Fig. 6i-j**). Consistent with large TEAD1 condensates serving as depots in TEAD1 condensate – positive cells (UOK121 and UOK342), their dissociation and thereby the release of TEAD1 to the nucleoplasm under hyperosmotic stress were accompanied by an increase in TEAD1 target gene expression, while this effect was not observed in TEAD1 condensate – negative cells (HK2 and UOK122, **Fig. 6k-m**). In summary, we have found that large TEAD1 condensates are seeded specifically in the pericentromeric region by the clustered MCAT sequences, serving important roles to buffer downstream gene expression.

### TEAD1 condensates selectively incorporate protein components

To further understand the functions of TEAD1 condensates, we set out to identify protein components of TEAD1 condensates using proximity labeling mass spectrometry^57,58^ with UOK342 cells stably expressing miniTurbo-TEAD1-HaloTag (**Fig. 7a**): miniTurbo (mT) is a compact biotin ligase for rapid labeling of TEAD1-surrounding proteins with biotin^57^, and HaloTag is for visualizing TEAD1 localization with Halo dyes. We used two additional cell lines as controls: UOK342 cells expressing miniTurbo-TEAD1 H3Mut-HaloTag, which form no condensates, and UOK342 cells expressing miniTurbo-HaloTag-NLS which localizes to the nucleus diffusely using a nuclear localization sequence (NLS, **Fig. 7a, b**). We first used imaging to confirm the efficient biotinylation with mT after addition of exogenous biotin in all three cell lines (**Fig. 7b**). We then enriched the biotinylated proteins from the whole cell lysate using streptavidin beads and confirmed that protein biotinylation increased upon biotin addition and all three constructs were expressed at similar levels (**Extended Data Fig. 8a, b**). Using mass spectrometry with tandem mass tag (TMT) labeling, we identified a total of 7580 proteins in WT, H3Mut and NLS samples combined (FDR < 0.01, **Supplementary table. 4**). Comparing the combined dataset of WT and H3Mut TEAD1 with the NLS control, we first identified the non-specific nuclear proteins (**Fig. 7c, labeled in blue**). Then, we compared the differential enrichment of proteins between WT TEAD1 and H3Mut group and identified 186 proteins enriched in WT TEAD1 and 409 proteins enriched in H3Mut TEAD1 (**Fig. 7d**). Among the 186 proteins enriched in the WT TEAD1 sample, 0% were non-specific nuclear proteins identified in Fig. 7c, while this percentage increased dramatically to 56% in the proteins enriched in the H3Mut TEAD1 sample, demonstrating the specificity of our mass spectrometry approach for detecting TEAD1 condensate components (**Fig. 7e**). GO terms of the unique proteins in WT TEAD1 sample revealed nuclear proteins associated with DNA binding and transcriptional regulation, consistent with TEAD1’s nuclear functions (**Fig. 7f, Extended Data Fig. 8c**). Using Co-IF, we confirmed the colocalization of endogenous RPA2 (replication protein A2, part of the trimeric RPA complex for maintaining genome stability) with TEAD1 condensates in UOK342 (**Fig. 7g, h**). RPA2 is not essential for TEAD1 condensate formation, as knocking it down did not affect TEAD1 condensate number or size (**Fig. 7i, j, Extended Data Fig. 8d-g**). Instead, RPA2 may be a client that partitions into TEAD1 condensates, as the fluorescence of RPA2 recovered much faster than TEAD1 after photobleaching (**Fig. 7k**)^59,60^. These results indicate that in addition to storing TEAD1, TEAD1 condensates may selectively incorporate proteins to maintain chromatin structure.

**Figure 7.**
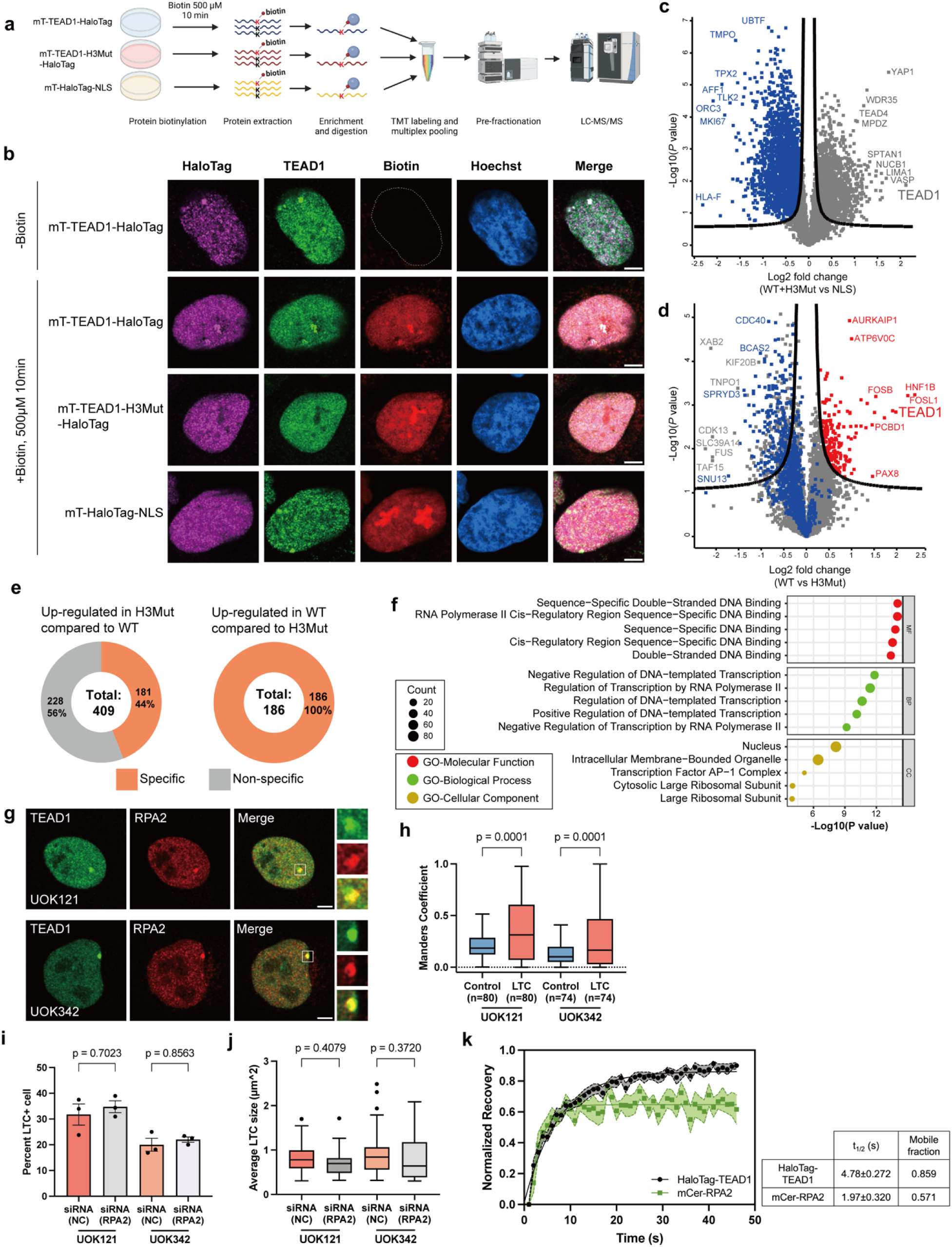
Identification of protein components of TEAD1 condensates with proximity-based mass spectrometry. **(a)** Schematic overview of the proximity labeling proteomic workflow used to identify TEAD1-associated proteins. Cells expressing miniTurbo-TEAD1 WT-, H3Mut-, or NLS-HaloTag constructs were incubated with biotin for 10 min to induce biotinylation, followed by protein extraction, streptavidin-based enrichment, on-bead digestion, and TMT labeling and pooling. The multiplexed samples were fractionated and analyzed by LC–MS/MS to compare specific TEAD1-dependent and non-specific biotinylation profiles. **(b)** Representative immunofluorescence images of UOK342 stably expressing miniTurbo-TEAD1 WT-, H3Mut-, or NLS-HaloTag with or without biotin treatment, biotinylation (red) in relation to HaloTag (magenta), TEAD1 (green), and nuclei (blue). Scale bars represent 5 μm. **(c)** Volcano plot of the biotinylated proteins identified from cells expressing miniTurbo-TEAD1 WT and H3Mut constructs, together with the NLS control representing nonspecific biotinylation. The x-axis represents the fold change comparing the combined TEAD1 WT and H3Mut conditions against the NLS control on a Log₂ scale, and the y-axis represents the p-value of the statistical analysis on a –Log₁₀ scale. The curved line indicates the boundary corresponding to a q-value of 0.05. Proteins with q < 0.05 and upregulated in the NLS condition shown in blue. **(d)** Volcano plot comparing biotinylated proteins between cells expressing miniTurbo-TEAD1 WT and H3Mut constructs. The x-axis represents the Log₂ fold change (WT vs H3Mut), and the y-axis represents the –Log₁₀(p-value). The curved line indicates the threshold corresponding to a q-value of 0.05. Red symbols (two-sided t-test, q < 0.05) denote proteins significantly enriched in the TEAD1 WT condition, whereas blue symbols indicate proteins belonging to the nonspecific biotinylation group identified in **(c)**. **(e)** Evaluation of nonspecific protein biotinylation in H3Mut and WT conditions. Among the significantly enriched proteins with q-value less than 0.05, nonspecific binding resulting from nonspecific protein biotinylation was evaluated. Donut charts show the number and proportion of proteins upregulated in H3Mut and WT, respectively. Proteins were categorized as specific or nonspecific based on nonspecific biotinylation, and inner labels indicate the total number of proteins and the percentage of each category. **(f)** Gene Ontology enrichment analysis of proteins significantly enriched under the TEAD1 WT condition with q value less than 0.05, as shown in red in (C). Enriched GO terms were categorized into molecular function in red, biological process in green, and cellular component in yellow. The top five significantly enriched GO terms for each category are displayed based on P values, and the circle size represents the number of proteins enriched in each term. **(g)** Representative Immunofluorescent images of endogenous RPA2 and TEAD1 in UOK121 and UOK342. Scale bars represent 5 μm. Insets show magnified images of boxed regions. **(h)** Quantification of co-localization between RPA2 and TEAD1 at LTC shown in **(g)**. **(i-j)** Quantification of percentage LTC-positive cells **(i)** and LTC area **(j)** upon RPA2 Knock-down in UOK121 and UOK342. **(k)** Quantification of recovered fluorescence intensity of TEAD1 and RPA2 at LTC in UOK342 HaloTag-TEAD1 Knock-In cell line. Shaded areas indicate the standard error of the mean (SEM) for each measurement (n = 11). Statistical significance determined by one-way ANOVA.

## Discussion

In this study, we made an interesting discovery that TEAD1 forms two types of condensates that differ in both size and function: small, transcriptionally active TEAD1 condensates that are the canonical condensates that colocalize with YAP and the large, transcriptionally silent TEAD1 condensates that localize to heterochromatin and function as depots to store excess TEAD1 protein at a relatively harmless genomic location. Cells need to maintain optimal concentrations of proteins for homeostasis. Too little protein may delay entry into mitosis^61,62^, and too much may promote protein aggregation and/or hyperactivation of downstream signaling^63,64^. Additionally, aberrant activation of oncogenes in non-transformed cells often induces oncogene-induced senescence involving cell cycle arrest^65–67^. Multiple cellular processes contribute to protein homeostasis including regulation of transcription, translation, and protein degradation systems. We propose that protein condensation of TEAD1 represents another way to regulate TEAD1 concentration and is used to both buffer and to limit the damaging effects caused by excess TEAD1 that may confer survival advantages to RCC cells. In RCC cells, YAP/TAZ and TEAD1 can activate genes mediating ferroptosis, an iron-dependent cell death pathway^68,69^. Preventing excessive TEAD1 activity by forming inactive TEAD1 condensates could represent a novel way to protect cancer cells from ferroptosis. On the other hand, not all phenomena in cancer cells promote cancer. TEAD1 condensates may instead represent a futile attempt of these cells to prevent tumorigenesis by decreasing YAP/TEAD-mediated cell proliferation. While we have successfully used hyperosmotic stress to disrupt TEAD1 condensates, future studies will investigate physiological modulation of TEAD1 condensates to examine their roles in cellular fitness and cancer progression.

Intriguingly, large TEAD1 condensates do not form at random locations in the nucleus, but instead localize to pericentromeric heterochromatin regions, long considered a transcriptionally inactive region critical for maintaining centromere integrity and proper segregation of chromosomes during mitosis^43,44^. While most transcription factors are absent from those areas, a few can be found there. Heat shock transcription factor 1 (HSF1) binds to the pericentromeric region of the human chromosome 9 during heat shock, transcribes satellite III repeats, and forms nuclear stress granules containing HSF1^70,71^. These HSF1 granules are thought to sequester active transcription complexes, such as mammalian CREB-binding protein, to shut down global transcription during heat shock^72^. Another transcription factor that localizes to the pericentromeric regions is GAGA factor, which binds to the GA/CT repeats and establishes chromatin boundaries in cooperation with Polycomb proteins and the inner nuclear membrane^73,74^. Ikaros is a protein that localizes to the centromeric heterochromatin of lymphocytes, forms condensates, and represses nearby genes^75^. Here we propose a novel role of the pericentromeric region, which is to serve as a reservoir for excess TEAD1 in a DNA-binding dependent manner. Indeed, when we overexpressed TEAD1 in other condensates-negative cell lines, we observed that TEAD1 condensates formed at heterochromatin and were likely to be pericentromeric (**Extended Data Fig. 5h, l, m**), indicating that this is a general mechanism utilized by cells to store TEAD1. This raises the question of whether other transcription factors can also localize specifically to pericentromeric or other heterochromatic regions and form condensates, and if so, what are the functions of these condensates?

Recently, condensate-targeting therapies have emerged as a promising way to treat previously untreatable diseases such as neurodegeneration and cancer^15,76–78^. Our data on TEAD1 condensates in RCC reveal that not all the condensates observed in diseases may drive that disease. In addition, a single transcription factor can also form condensates of different sizes and functions in different cell types^15^. Therefore, careful studies of condensate morphology and function are paramount for designing precise condensate-targeting treatments.

## Supporting information

Supplementary Table 1_Primers

Supplementary Table 2_Antibodies

Supplementary Table 3_DNA Oligos

## Acknowledgements

We appreciate the discussions and feedback from Cai lab members, Sam Botterbusch and Anthony Leung, the centromere DNA-FISH probe gifted by Alan Meeker, and the assistance from Johns Hopkins School of Medicine Single Cell & Transcriptomics Core and Center for Proteomics Discovery. This work is supported by the Department of Defense Kidney Cancer Idea Development Award (W81XWH2210900, D.C.) National Institutes of Health (T32GM080189, K.S.L.; T32CA009110, J.D., E.L., E.B.; R35GM56573, J.M.K.; R35GM142837, D.C.; R35GM133712, C.Z.; and intramural grant, W.M.L.), Johns Hopkins University Catalyst Award (D.C.), National Science Foundation Graduate Research Fellowship Program (DGE-1745301, S.R.Y), the Pew-Stewart Scholar for Cancer Research Award (S.C.), the Searle Scholar Award (S.C.), the Merkin Innovation Seed Grant (S.C.), the Mallinckrodt Research Grant (S.C.), the Margaret E. Early Medical Research Trust Grants (S.C.), and the Alex’s Lemonade Stand Foundation Innovation Grant (No. 1260879, S.C.).

---

**Extended Data Fig. 1.**
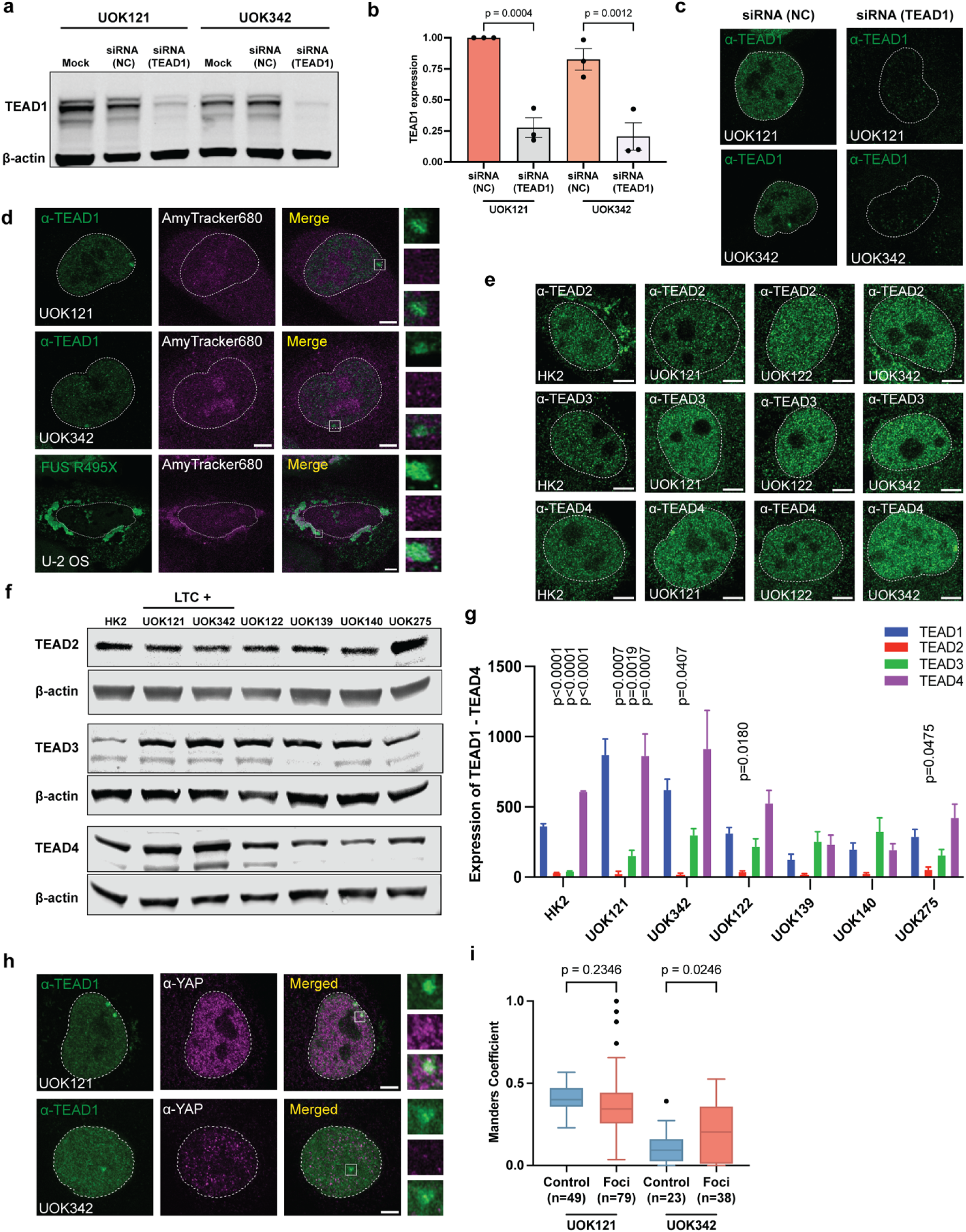
TEAD1 condensates are not protein aggregates or artifacts. **(a-b)** Immunoblotting of efficient TEAD1 knockdown in UOK121 and UOK342 cells in **(a),** Mock: no transfection. Quantification of TEAD1 expression between TEAD1 siRNA treatment and negative control (NC) in UOK121 and UOK342 in **(b)**, data are shown as mean ± SEM. **(c)** Immunofluorescence images of TEAD1 (green) in UOK121 and UOK342 cells upon treatment with TEAD1 knockdown siRNA (TEAD1) or control siRNA (NC). Scale bars represent 5 μm. Nuclear boundaries are indicated by dotted lines. **(d)** Representative 2D immunofluorescence images showing AmyTrakcer680 staining do not overlap with TEAD1 condensates visualized by TEAD1 antibody, but overlaps with protein aggregates formed by expressing FUS R495X mutant^79^ (detected by Myc antibody) in U-2 OS cells. Scale bars are 5μm. Insets show magnified images of boxed regions. Nuclear boundaries are indicated by dotted lines. **(e)** Representative 2D immunofluorescence images showing TEAD2-4 staining in HK2, UOK121, UOK342, and UOK122 cells. Scale bars are 5μm. Nuclear boundaries are indicated by dotted lines. **(f-g)** Immunoblotting of TEAD2-4 in LTC-positive and LTC-negative cell lines in **(f)**. Quantification of TEAD1-4 expression band intensity cross-compared in HK2 and UOK cells in **(g)**, data normalized to β-actin. Data are presented as mean ± SEM. Statistical significance was determined by one-way ANOVA, only p-values <0.05 are shown. **(h)** Representative immunofluorescence images of TEAD1 condensates and their spatial relationship with YAP in UOK121 and UOK342. Insets show magnified images of boxed regions. Scale bars are 5μm. Nuclear boundaries are indicated by dotted lines. **(i)** Quantitative analysis and statistical evaluation of TEAD1 condensates co-localization with YAP. Co-localization was assessed using Manders’ coefficient within defined regions surrounding TEAD1 condensates, with control measurements taken from random nucleoplasmic regions. Statistical significance was determined by one-way ANOVA.

**Extended Data Fig. 2.**
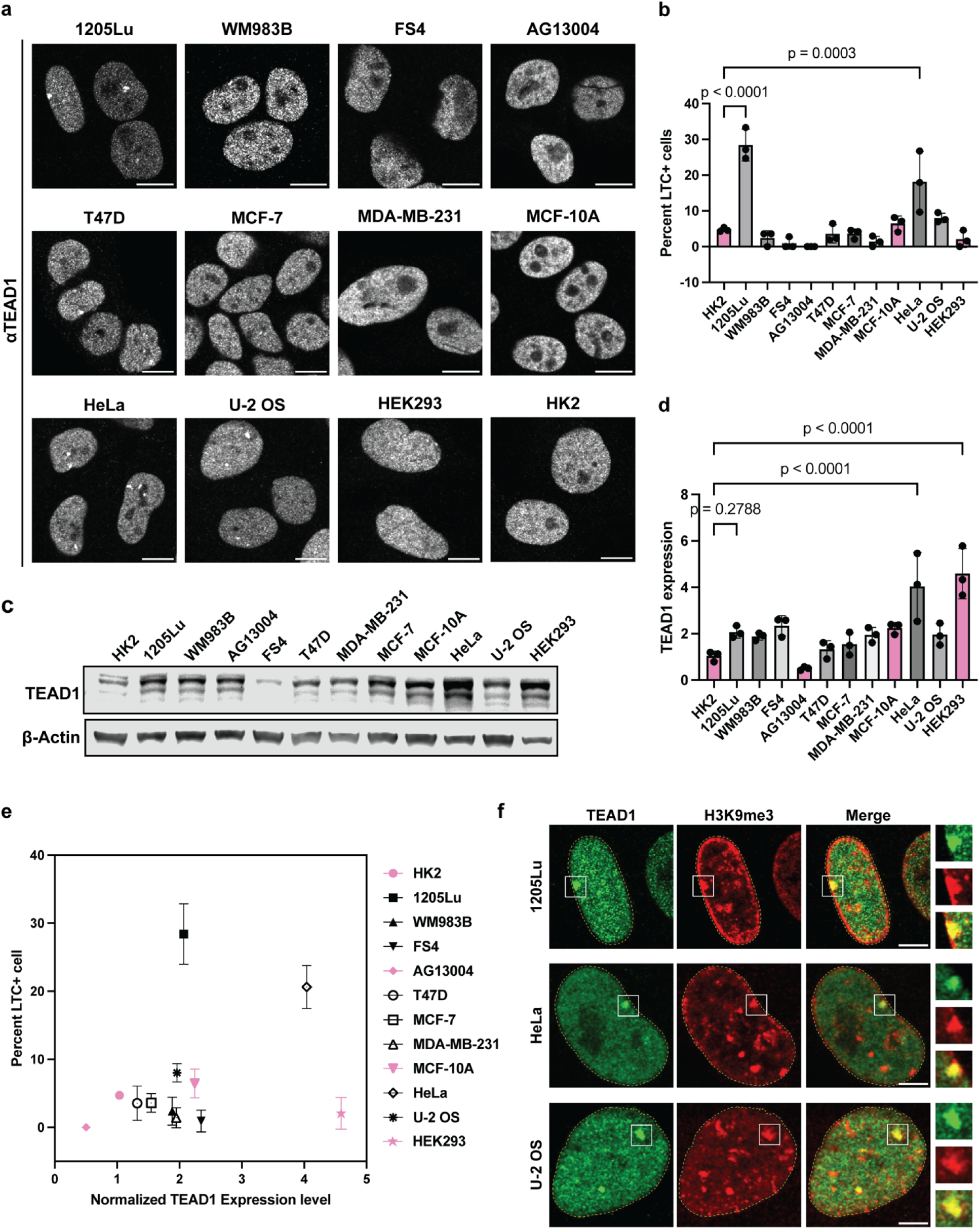
TEAD1 forms condensates in other cancer cell lines. **(a)** Representative immunofluorescence images of TEAD1 staining in different cancer (Melanoma: 1205Lu, WM983B, FS4; Breast cancer: T47D, MCF-7, MDA-MB-231; Cervical cancer: HeLa; Osteosarcoma: U-2 OS) and non-cancer cell lines (Fibroblast: AG13004; mammary gland epithelial cells: MCF-10A; embryonic kidney cells: HEK293). Scale bars = 5µm. **(b)** Quantification of percent LTC-positive cells in various cell lines shown in **(a)**. **(c-d)** Immunoblot of TEAD1 expression in cell lines shown in **(a)**, quantification and statistical analysis of the expression bands in **(d)**. Data normalized to β-actin and TEAD1 expression in HK2. **(k)** Scatter plot of TEAD1 protein expression by percent LTC-positive cells in cancer cell lines (black) and non-cancer cell lines (pink). **(l)** Representative immunofluorescent images of TEAD1 (green) and H3K9me3 (Red) in 1205Lu, Hela, and U-2 OS cell lines. Scale bars = 5µm. Insets show magnified images of boxed regions. Nuclear boundaries are indicated by dotted lines.

**Extended Data Fig. 3.**
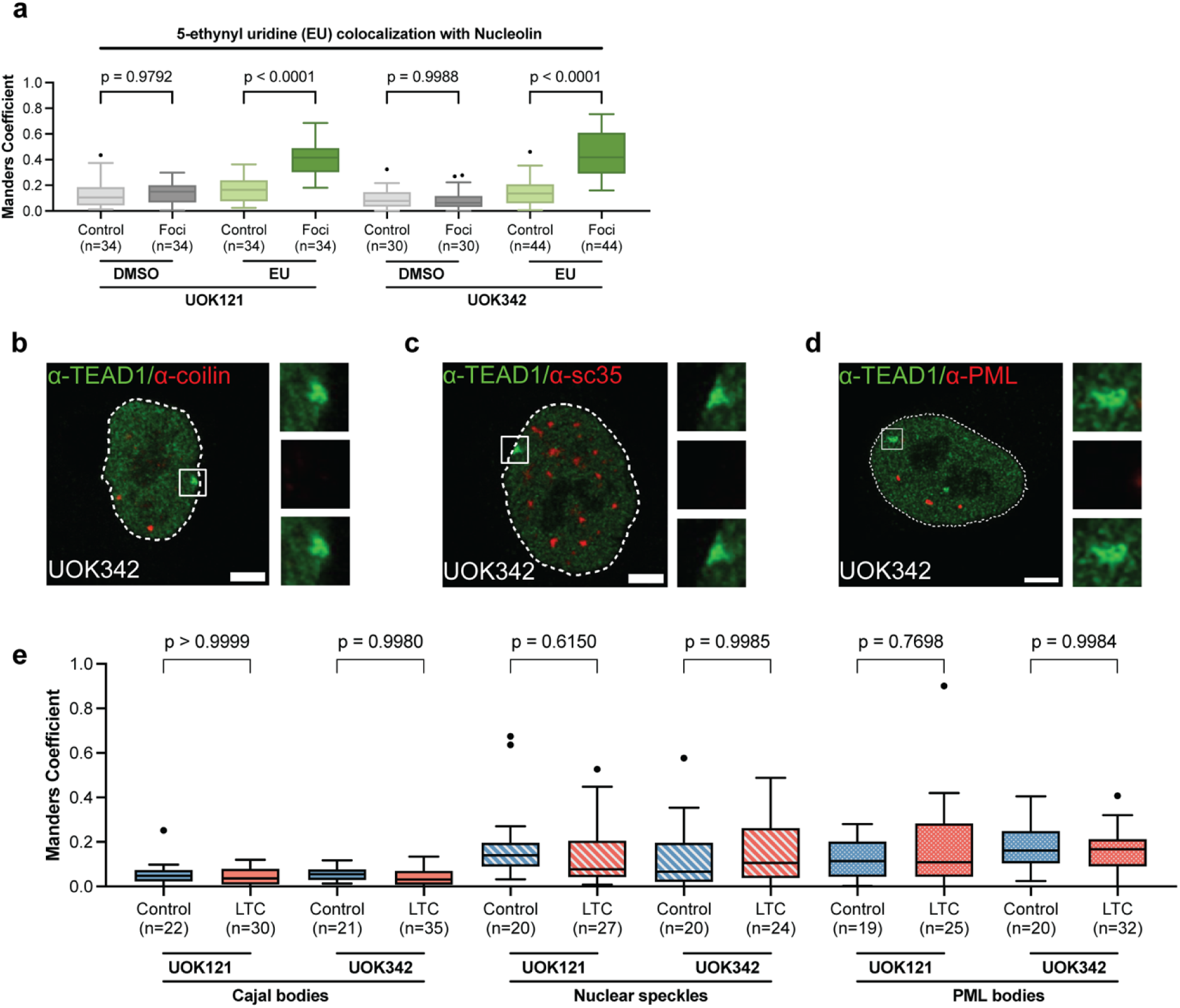
TEAD1 condensates do not colocalize with other nuclear bodies. **(a)** Quantification of colocalization of nucleolin with EU in UOK121 and UOK342 treated with DMSO control or EU for 10min, measurement made around nucleolin condensates with comparable size of LTC and control regions, as shown in **Fig.2c**. **(b-d)** Representative UOK342 immunofluorescence images of TEAD1 condensates (green) with various Nuclear Bodies (red): Cajal bodies **(b)**, nuclear speckles **(c)**, and PML bodies **(d)**. Scale bars are 5μm. Nuclear boundaries are indicated by dotted lines. **(e)** Quantification of co-localization of immunofluorescence imaging of TEAD1 with indicated MLO markers in UOK121 and UOK342 around LTC regions and randomly selected nuclear regions (control) in **(b-d)**. Statistical significance determined by one-way ANOVA.

**Extended Data Fig. 4.**
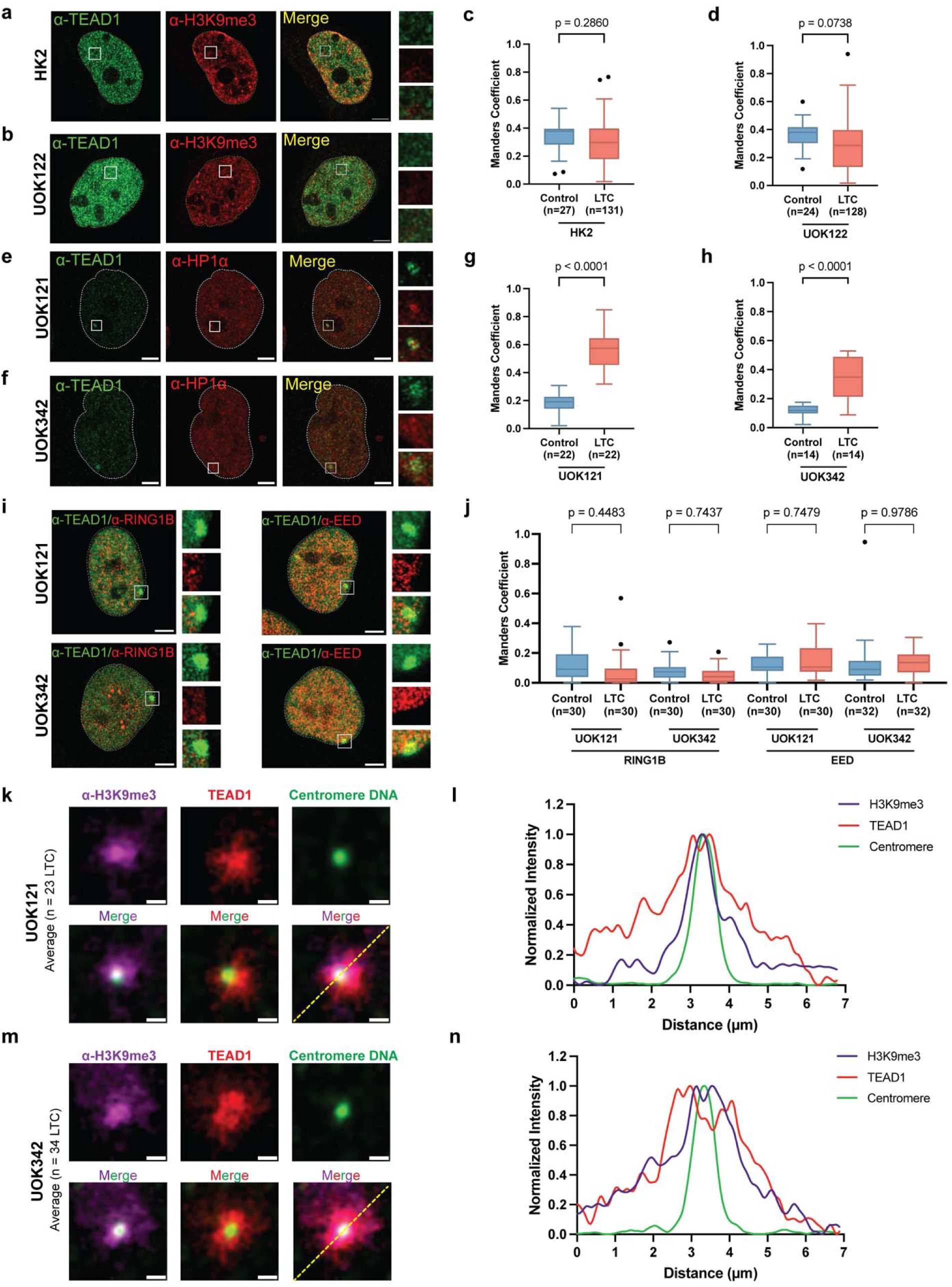
TEAD1 condensates colocalize with heterochromatin markers but not the polycomb bodies. **(a-d)** Representative immunofluorescence images of TEAD1 (green) and H3K9me3 (red) in HK2 **(a)** and UOK122 **(b)** cells, scale bars = 5 μm. Insets show magnified images of boxed regions. Co-localization analysis between TEAD1 and H3K9me3 around LTC and control nuclear regions (control) in HK2 **(c)** and UOK122 **(d)** cells. **(e-h)** Representative immunofluorescence images illustrate the co-localization of TEAD1 (green) with HP1 (red) in UOK121 **(e)** and UOK342 **(f)** cells. Insets show magnified images of boxed regions. Co-localization analysis between TEAD1 and HP1 around LTC and control nuclear regions (control) in UOK121 **(g)** and UOK342 **(h)** cells. Statistical significance determined by unpaired t-test. **(i-j)** Representative immunofluorescence images illustrate the co-localization of TEAD1 (green) with Polycomb Repressive Complex 1 (PRC1) marker RING1B or Polycomb Repressive Complex 2 (PRC2) marker EED (Red) in UOK121 and UOK342. Insets show magnified images of boxed regions. Co-localization analysis between TEAD1 and Polycombs around LTC and control nuclear regions (control) in UOK121 and UOK342 in **(j)**. Statistical significance determined by one-way ANOVA. **(k-n)** Average intensities of images centered at the centromere in UOK121 **(k)** and UOK342 **(l)**. UOK121 n=23 condensates, UOK342 n=23 condensates. Line plots of H3K9me3 (magenta line), TEAD1 (red line), and centromere DNA-FISH signal (green line) in UOK121 **(m)** and UOK342 **(n)**.

**Extended Data Fig. 5.**
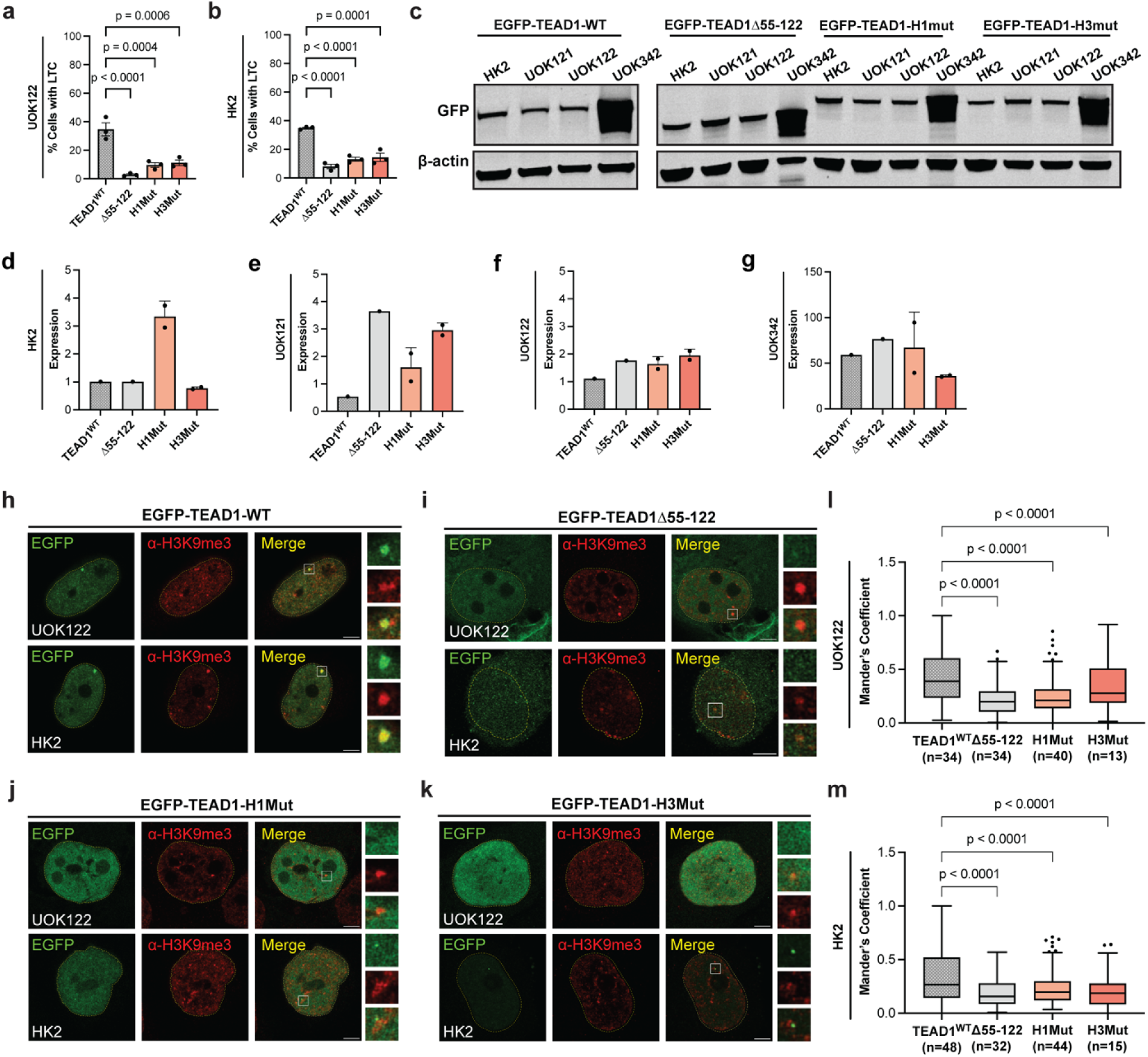
TEAD1 condensates formation is mediated by its DNA-binding domain. **(a-b)** Quantification of percentage LTC-positive cells upon transfection in UOK122 **(a)**, and HK2 **(b)**. Data are presented as mean ± SEM. **(c)** Immunoblot shows EGFP expression level after transfection in HK2 and UOK cell lines. **(d-g)** Quantification of immunoblot proteins expression levels in **(a)** with GFP antibodies across different indicated cell lines. The p values are determined by unpaired t test comparing EGFP-TEAD1-WT and EGFP-TEAD1-H3mut expression levels. **(h-k)** Representative images of UOK122 and HK2 cells expressing EGFP-TEAD1-WT **(h)**, EGFP-TEAD1-DNA BD deletion **(i)**, EGFP-TEAD1-H1Mut **(j)**, and EGFP-TEAD1-H3Mut **(k)**, with immunofluorescence signal of H3K9me3 (red). Insets show magnified images of boxed regions. Scale bars = 5μm. **(l-m)** Quantification of co-localization between H3K9me3 and EGFP-TEAD1 variants shown in **(h-k).** Statistical significance determined by one-way ANOVA.

**Extended Data Fig. 6.**
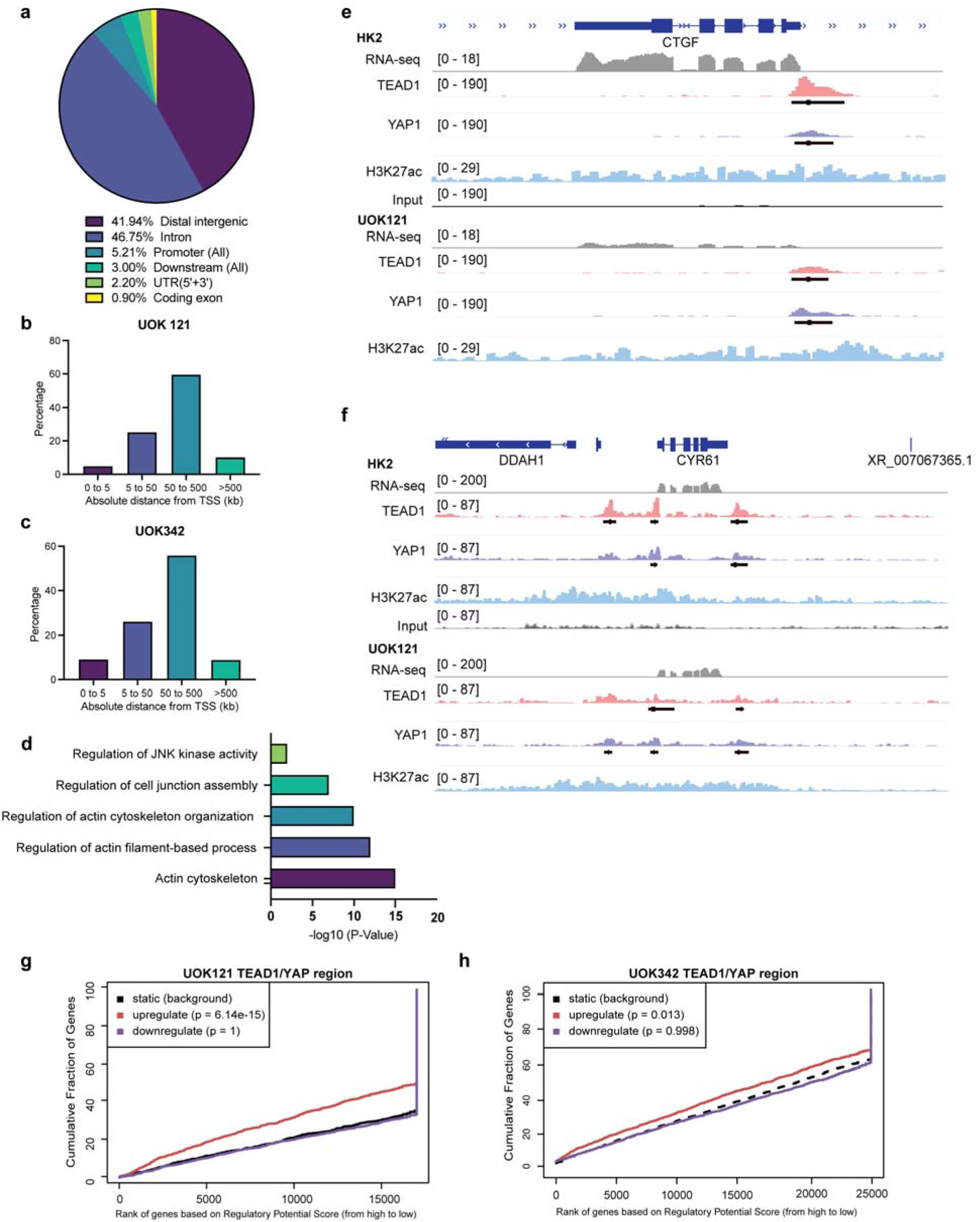
TEAD1 binds to both enhancers and heterochromatin regions in the genome. **(a)** TEAD1 peaks were mapped back to various regions of the genome. The percentage of the TEAD1 peaks found within each region of the genome is shown in the pie chart. **(b-c)** Bar graph from GREAT analysis showing the percent distribution of TEAD1/YAP peaks in relation to the transcription start site (TSS) of genes in UOK121 **(b)** and UOK342 **(c)** cells. Distance to TSS is shown in kilobases (kb). **(d)** Gene ontology analysis of TEAD1-bound peaks using DAVID under default parameter. **(e-f)** Gene tracks showing TEAD1/YAP1 binding at regulated genes CTGF **(e)** and Cyr61 **(f)**, with their corresponding mRNA expression profiles in HK2 and UOK121 cells. **(g-h)** Line graph showing correlation of TEAD1 ChIP-seq binding data with differential gene expression in UOK121 **(g)** and UOK342 **(h)** using Binding and Expression Target Analysis (BETA). Genes were ranked by regulatory potential score (x-axis) based on ChIP-seq peaks that are co-bound by TEAD1 and YAP. Cumulative fractions of upregulated (red), downregulated (purple), and static (black) genes were plotted (y-axis).

**Extended Data Fig. 7.**
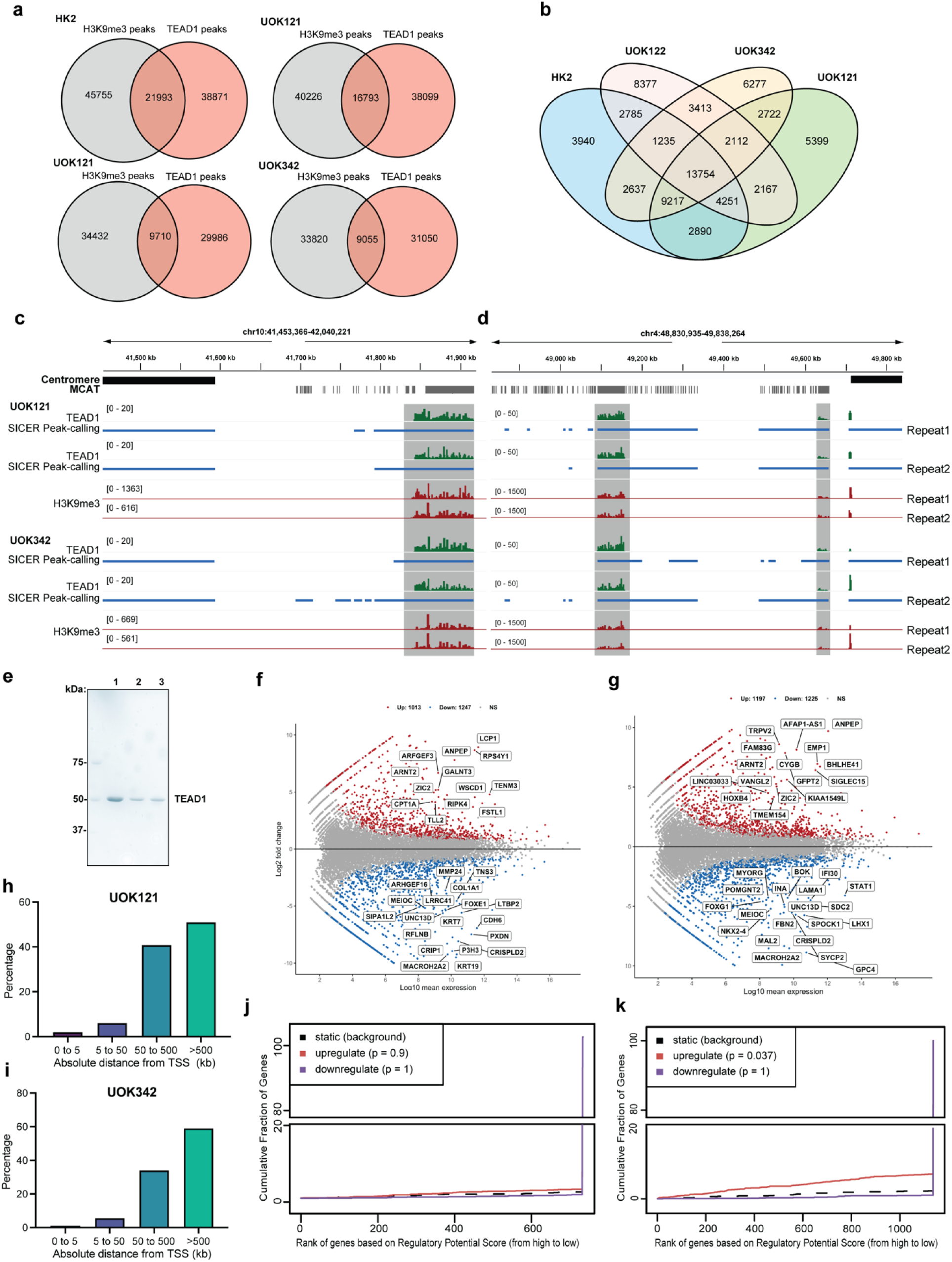
ChIP-seq reveals universal binding sites of TEAD1 condensates. **(a)** Venn diagram representing uniquely mapped TEAD1 peaks in HK2 and UOK cell lines, overlapped regions of H3K9me3. **(b)** Venn diagram representing shared TEAD1 overlapped regions of H3K9me3 across cell lines. **(c-d)** Gene tracks showing TEAD1/H3K9me3 in UOK121 and UOK342 cells at clusterd MCAT motif at pericentromeric region in chromosome 1 **(c)** and chromosome 4 **(d)**. The grey regions are showing intersected SICER peak-called TEAD1 and H3K9me3 region. TEAD1 is mapped with multimap parameter and then normalized with counts per million (CPM). TEAD1 tracks are shown in green and H3K9me3 tracks are in red. Two biological repeats are shown. **(e)** Purified TEAD1 for in-vitro droplet assay shown in Figure 5j. Coomassie stained SDS-PAGE gel of final purified TEAD1 samples. Lane 1 is unlabeled TEAD1, lane 2 is Alexa Fluor 488-labeled TEAD1, and lane 3 is a 10:1 mixture of unlabeled: labeled TEAD1. **(f-g)** MA plots showing differential gene expression from RNA-seq analysis comparing UOK121 **(f)** and UOK342 **(g)** to HK2 cells. Gene expression is plotted as log□ fold change (y-axis) versus log□□ mean expression (x-axis). Red dots represent significantly upregulated genes and blue represent downregulated genes (p ≤ 0.01). Non-significant genes are shown in gray. **(h-i)** Bar graph from GREAT analysis showing the percent distribution of TEAD1/H3K9me3 peaks in relation to the transcription start site (TSS) of genes in UOK121 **(h)** and UOK342 **(i)** cells. Distance to TSS is shown in kilobases (kb). **(j-k)** Line graph showing correlation of TEAD1 ChIP-seq binding data with differential gene expression in UOK121 **(j)** and UOK342 **(k)** using BETA. Genes were ranked by regulatory potential score (x-axis) based on ChIP-seq peaks that are co-bound by TEAD1 and H3K9me3. Cumulative fractions of upregulated (red), downregulated (purple), and static (black) genes were plotted (y-axis).

**Extended Data Fig. 8.**
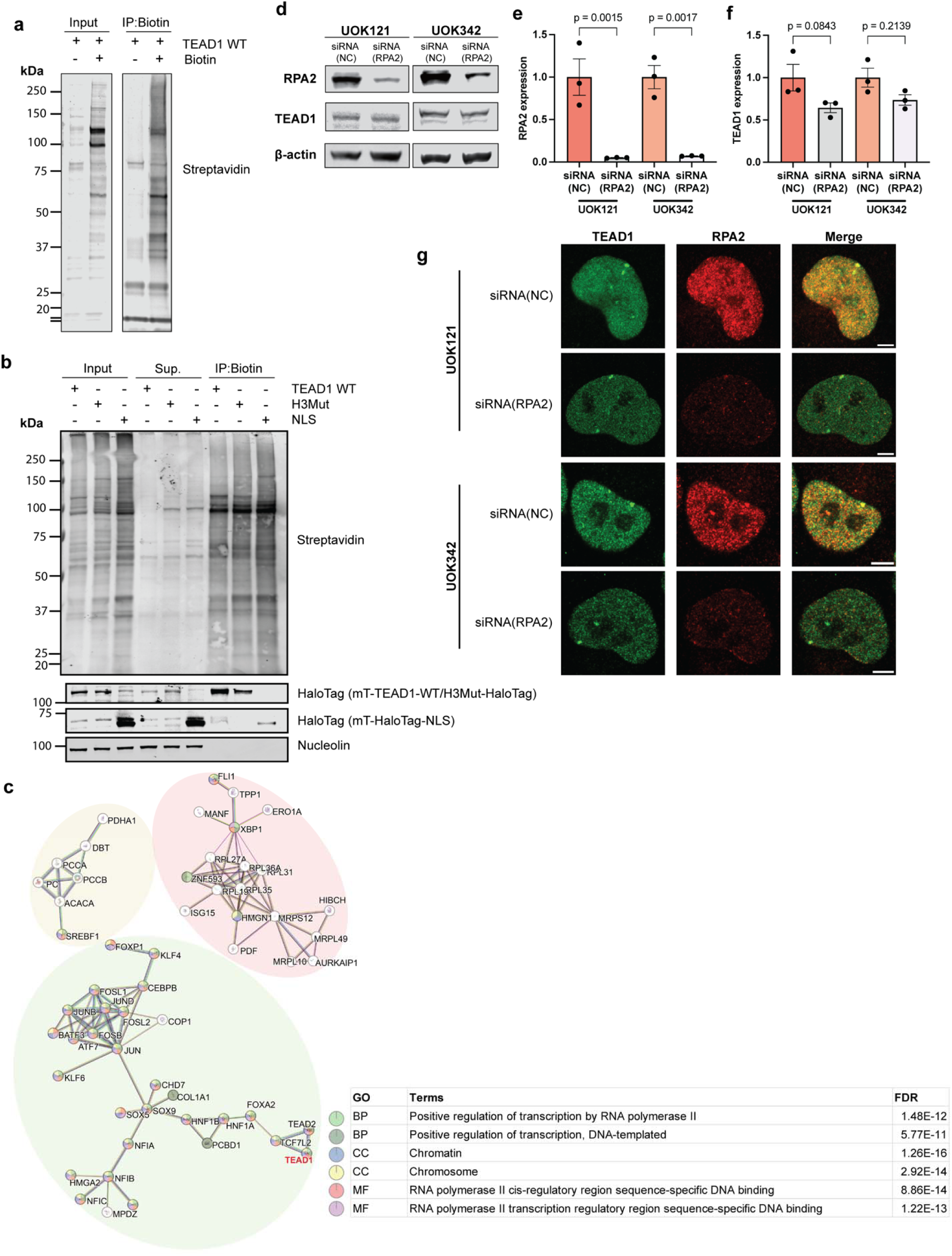
Identification and validation of protein components in TEAD1 condensate. **(a)** Western blot showing enriched biotinylated proteins (Streptavidin IRDye680) with or without biotin treatment of UOK342 cells expressing miniTurbo-TEAD1-HaloTag. **(b)** Western blot comparing biotinylated protein enrichments of miniTurbo-TEAD1 WT-, H3Mut-, and NLS-HaloTag stably expressed in UOK342 cells upon biotin treatment. **(c)** Interactome analysis of proteins significantly enriched in the TEAD1 WT condition (red group) using STRING protein–protein interaction (PPI) analysis. The analysis was performed with high confidence (interaction score threshold of 0.7). The resulting network contained 56 nodes and 105 edges, with an average node degree of 1.47 and an average local clustering coefficient of 0.381. The expected number of edges was 51, and the observed network showed significantly more interactions than expected (PPI enrichment p < 1.0 × 10□¹□). Clusters composed of fewer than four proteins were removed for visualization. TEAD1 is highlighted in red. Protein–protein interaction analysis was conducted using STRING based on the evidence types of text mining (yellow edges), experiments (purple edges), databases (blue edges), and co-expression (black edges). **(d-f)** Immunoblotting of efficient RPA2 knockdown in UOK121 and UOK342 cells in **(d).** Quantification of RPA2 expression **(e)** and TEAD1 expression **(f)** between TEAD1 siRNA treatment and negative control (NC) in UOK121 and UOK342, data are shown as mean ± SEM. Statistical significance determined by one-way ANOVA. **(g)** Immunofluorescence images of TEAD1 (green) and RPA2 (red) in UOK121 and UOK342 cells upon treatment with RPA2 knockdown siRNA (RPA2) or control siRNA (NC). Scale bars represent 5 μm.

## Materials and Methods

### Data availability statement

The accession to RNA seq data and ChIP seq data (FASTQ and BAM files) reported in this paper is SRA PRJNA1268000, and the processed data (BIGWIG) is at Gene Expression Omnibus GSE298316. The mass spectrometry proteomics data have been deposited to the ProteomeXchange Consortium via the PRIDE^80^ partner repository with the dataset identifier PXD072858 and 10.6019/PXD072858. Reviewer access token: Q270kaz74QYY. Data will be released upon publication. All other data supporting the findings of this study are available within the paper and its Supplementary Information.

### Availability of materials

The UOK cell lines are derived from National Cancer Institute on a Material Transfer Agreement therefore are not readily available. All other materials are promptly available upon request.

### Code availability

We do not report the development of new codes.

## Method

### Cell culture, transfection, and siRNA

Patient-derived immortalized cell lines UOK121, UOK139, UOK140, and UOK275 cells were gifted from W. Marston Linehan, MD, Center for Cancer Research, National Cancer Institute. HK-2 is from ATCC (CRL-2190). Cells were cultured at 37°C and 5% CO2 in DMEM supplemented with 10% fetal bovine serum (FBS; Gibco, 26140079), 100 U/mL (1%) penicillin/streptomycin (Gibco, 15140122), 2 mM (1%) GlutaMAX-l (Gibco, 35050061), 1x NEAA (Gibco, 11140050). Cell lines 1205Lu, WM983B, FS4^81^, and AG13004 (Coriell Institute) were gifted from Ashani Weeraratna, PhD, Biochemistry and Molecular Biology, Johns Hopkins Bloomberg School of Public Health. 1205Lu, WM983B, and AG13004 cells were cultured in DMEM (Gibco, 12430-047) supplemented with 10% FBS. FS4 cells were cultured in RPMI (Gibco, 11875-085) and supplemented with 10% FBS and 1% GlutaMAX (Gibco, 35050-061). T47D, MCF-7, MDA-MB-231, and MCF-10A cells were gifted from Brittany Jenkins-Lord, PhD, Biochemistry and Molecular Biology, Johns Hopkins Bloomberg School of Public Health. T47D cells were cultured in RPMI-1640 (ATCC, 30-2001) with 10% FBS (Gibco, A52558-01) and 10µg/ml insulin (Lonza, BE02-033E20); MCF-7 cells were cultured in EMEM (ATCC, 30-2003) supplemented with 10% FBS and 10µg/ml insulin; MDA-MB-231 cells were cultured in DMEM (ATCC, 30-2002) supplemented with 10% FBS; MCF-10A cells were cultured in DMEM-F12 (ATCC, 30-2006) supplemented with 5% horse serum (Millipore, H1138), 20ng.ml EGF (Rockland, 009-001-C26), 0.5mg/mL Hydrocortisone (VWR, 1315004), 100ng/mL Cholera toxin (Millipore, 9012-63-9), 10µg/ml insulin and 1% penicillin/streptomycin (VWR, 10220-718). For overexpression experiments UOK and HK2 cells were transfected with EGFP-TEAD1 wild-type, DNA binding mutants, and other EGFP-TEAD1 variants using Lipofectamine 3000 Transfection Reagent (Thermo Fisher, L3000015), for 16 h. For siTEAD1 experiments, TEAD1 siRNA (Thermo Fisher, Silencer Select s13962), RPA2 siRNA (Horizon Discovery, J-017058-09-0002) or scrambled negative control siRNA (Thermo Fisher, AM4611) was used at a final concentration of 10 nM, and was transfected into cells using the Lipofectamine RNAiMAX transfection reagent (Thermo Fisher, 13778075) for 48 h, after which cells were replated for Western blot and imaging.

## Cell lines generation

### UOK342 HaloTag-TEAD1 KI cell line generation

Endogenously tagged UOK342 Halo-Tag KI cells were engineered using CRISPR-Cas9 system. pTEAD1-gDNA targeting the N-terminus of TEAD1 was assembled into PX458 template plasmid (Addgene #48138). pTEAD1-donor was assembled into pUC57mini (Addgene #88845) using gblock containing HaloTag sequence and 300bp homology arms of the gDNA target site. UOK342 cells were transfected with lipofectamine 3000 (L3000015, ThermoFisher) following the manufacture’s protocol. The cells were cultured for >14 days. Cells were stained for 30 min with JF549-HaloTag ligand (100nM, Janelia Materials) and sorted by fluorescence-activated cell sorting (FACS) and expanded. For cell line validation, the genomic DNA from the selected clone was extracted using GeneJET Genomic DNA Purification Kit (K0721, Thermo Scientific). PCR product of edited region was amplified using Q5 High-Fidelity 2X 901 Master Mix (M0492, NEB) and verified using Sanger sequencing (See **Supplementary Table 1** for sequence).

### miniTurbo plasmid construct and stable cell line generation

miniTurbo-TEAD1-HaloTag, miniturbo-TEAD1 H3Mut-HaloTag, and miniTurbo-HaloTag-NLS expression plasmids were generated by high-fidelity of the PCR product amplified from EGFP-TEAD1 vector, and 3xHA-miniTurbo-NLS_pCDNA3 (Addgene, Plasmid #107172) (See **Supplementary Table 1** for sequence). Plasmids were validated by whole-plasmid sequencing (Plasmidsaurus). For cell line generation, miniturbo-TEAD1-HaloTag and miniturbo-TEAD1 H3Mut-HaloTag plasmids were transfected in UOK342 cells using Lipofectamine 3000 Transfection Reagent (L3000015, ThermoFisher) and passaged for >14 days. Cells were stained for 30 min with JF549-HaloTag ligand (100nM, Janelia Materials). Cells with stable expression were sorted by fluorescence-activated cell sorting (FACS) and expanded. Expression of the fusion proteins were validated using western blot.

### Immunofluorescence staining

After transfection, UOK and HK2 cells were plated on coverslips pre-coated with fibronectin (7.5 μg/mL; Millipore, FC010). Cells were grown for 16 hours and fixed with 4% FA (EMS), permeabilized with 0.1% Triton X-100, and blocked with 3% BSA in 1X PBS. The cells were then incubated overnight with primary antibodies in 1% BSA at 4°C and incubated with Alexa Fluor-conjugated secondary antibodies in 1% BSA for 1 hour at RT (**Supplementary Table 2**). For imaging and quantification, at least 20 fields of view per coverslip were randomly chosen by Hoechst nuclear staining and imaged using a Zeiss LSM900 Airyscan microscope, followed by Airyscan processing (2D, default settings).

### Domain deletion and site-directed mutagenesis

The plasmid EGFP-TEAD1 was first digested using BamHI and ApaI to create the backbone. This step was followed by the amplification of two specific DNA fragments from the parental EGFP-TEAD1 plasmid. To generate EGFP-TEAD1Δ55-122, Primer set TEAD1-1 used for the first fragment, and primer set TEAD1-2 for the second fragment (See **Supplementary Table 1** for sequence). To generate the EGFP-TEAD1-H3Mut variant, the process began with BamHI digestion of EGFP-TEAD1, followed by PCR to generate two segments using primer sets TEAD1-3 and TEAD1-4. To generate EGFP-TEAD1-H1mut, PCR amplification was performed to produce three distinct fragments. The first fragment utilized primers TEAD1-3f and TEAD1-5r, the second employed TEAD1-5f and TEAD1-4r, and the third fragment was a previously prepared backbone. These PCR fragments were subsequently assembled into the final construct through the HIFI assembly method. For the creation of EGFP-TEAD1ΔIDR, PCR amplification was performed using primer pairs TEAD1ΔIDR 5’ of the PCR product was phosphorylated using T4 Polynucleotide Kinase (NEB, M0201S), parental plasmids were digested using DpnI restriction enzyme (NEB, R0176S), then the PCR product was ligated into plasmids using T4 DNA ligase (NEB, M0202S). For generating the EGFP-TEAD1 Y406H, S102A and S102D mutant construct, PCR amplification was performed using the parental EGFP-TEAD1 plasmids and primer pairs Y406H, S102A, S102D, respectively (**Supplementary Table 1**).

### DNA Fluorescence *in-situ* hybridization (FISH) and immunofluorescence staining of TEAD1 condensates

UOK and HK2 cells were plated on #1.5H high-performance coverslips (Marienfeld Superior) pre-coated with fibronectin (7.5 μg/mL; Millipore, FC010). Cells were grown overnight, washed with 1X PBS and fixed with 4% methanol-free formaldehyde (Cell Signaling 12606)/1X PBS) for 10 minutes. Coverslips were washed in 1X PBS / 0.002% Tween (PBS-T) 5 times with a transfer method detailed in ^82^, and permeabilized with 0.2% Triton-X-100 for 10 minutes before blocking in MaxBlock (Active Motif 15252) for 1 hour. Coverslips were incubated overnight in primary antibody (H3K9me3, Active Motif 39161 1:500; TEAD1, BD 610922 1:200) in MaxBlock, and washed 5X in PBS-T. Secondary antibodies (AF647 1:500; AF594 1:1000) were incubated in MaxBlock for 1 hour in the dark, and washed 5X in PBS-T before post-fixation in 4% methanol-free formaldehyde for 10 minutes. Subsequently, FISH hybridization for centromere probes was performed as described^83^. AF488-labelled probes (gift from Alan Meeker) suspended in hybridization buffer were denatured at 84°C for 5 minutes before hybridization at room temperature for 2 hours. Coverslips were then washed 2X in PNA buffer for 15 minutes at room temperature, then washed 2X in PBS-T and counterstained with Hoescht 33342 at 1:5000, before washing 2X in PBS-T, once in ddH2O, pre-equilibrating and mounting in Vectashield before proceeding to Airyscan imaging.

### Intron RNA FISH combined with immunofluorescence

Human *Cyr61*_intron with Quasar 570 dye (Biosearch Technologies, ISMF-2066-5), human *ACTB*_intron with Quasar 570 dye (Biosearch Technologies, ISMF-2002-5), Stellaris RNA FISH Hybridization Buffer (Biosearch Technologies, SMF-HB1-10), Stellaris RNA FISH Wash Buffer A (Biosearch Technologies, SMF-WA1-60) and Wash Buffer B (Biosearch Technologies, SMF-WB1-20) are purchased from Biosearch Technologies. We followed the protocol for sequential IF + FISH in Adherent Cells listed on the Biosearch Technologies website under Stellaris RNA FISH protocols.

### Nascent RNA labeling assay

UOK121 and UOK342 cells were cultured on coverslips and grown for 24 hours. Cells were treated with 1mM 5-ethynyl uridine (EU) or DMSO vehicle control for 10min. Nascent RNA labeling assay was conducted using Click-It RNA Alexa Fluor 594 Imaging Kit (Invitrogen, C10330) following the manufacturer’s protocol. Immunofluorescent experiments against TEAD1 and Nucleolin were conducted after cells washed with Click-It rinse buffer. Images were captured with Zeiss LSM900 Airyscan microscope and colocalization analysis was quantified using Image J software.

## Image analyses

### TEAD1 condensates nuclear localization quantification

Immunofluorescence staining against of UOK121 and UOK342 cell lines was performed. 3D images were acquired using Zeiss LSM 900 with Airyscan 2 detector. Z-stacks were captured at 0.15 µm intervals and 3D Airyscan processing was applied. TEAD1 condensates localization was analyzed in 3D using Zeiss Zen software. Nucleoli were identified based on nucleolin staining, and perinuclear regions were defined as areas within 1 µm of the nuclear envelope, as delineated by Hoechst staining. TEAD1 condensates were considered colocalized with the nucleolus if they overlapped with nucleolin-positive regions in 3D space. Similarly, TEAD1 condensates were classified as perinuclear if they were located within 1 µm of the nuclear envelope. Colocalization was visually confirmed in all three dimensions. The percentage of TEAD1 condensates localized to the nucleolus, perinuclear regions, both, or neither was calculated for each cell line. For representative images, maximum projections of the Z-stack were generated in Fiji.

### TEAD1 condensate quantification

Images were taken of fixed cells stained with Hoechst dye and TEAD antibody. Cellprofiler (Cellprofiler version 4.2.8) was used to identify both cell count and the appearance of TEAD condensates. Using the Hoechst as a nuclear marker we used Cellprofiler to identify the nuclei as objects. Using the TEAD antibody Cellprofiler thresholded the TEAD condensates based on their size and fluorescent intensity. The objects were manually screened for false positives, and any object that had been thresholded inaccurately was removed from the data set before the final quantification. To determine appearance rate of TEAD condensates we divided the number of cells who positively expressed at least 1 TEAD condensate by the total number of nuclei detected.

### Co-localization analysis

A square region measuring 3μm by 3μm was delineated around the focal point of TEAD1 condensates. Additionally, a negative control square was randomly selected within the nucleoplasm where TEAD1 staining appeared diffused. The colocalization analysis was conducted using FIJI software^84^ with BIOP-JACoP plugin^85^. Statistical evaluation was performed using an unpaired T-test in GraphPad prism. To calculate the enrichment of TEAD1 and YAP signal at *CYR61* Intron-RNA-FISH, we used a 2.12 μm x 2.12 μm region of interest (ROI) box to center around RNA FISH signal in 2D immunofluorescence images using ImageJ, and duplicated both TEAD, YAP and *CYR61* channels. The duplicated smaller images are then combined into stacks of each channel, and the averaged images are derived using the Z-project-> Average function. Enrichment of averaged signals across the center point were generated using the Plot Profile function in ImageJ. Similar analyses were done except for centering at the LTC or Centromere DNA FISH Signal.

### 3D Distance Analysis

Immunofluorescence staining of HK2 and UOK cell lines was performed. The imaging was conducted using Airyscan Z-stacks with a z-interval of 0.13 μm. Three-dimensional distance analysis of TEAD1 condensates with CENP-A or TRF2 was performed using ImageJ plugin Distance Analysis (DiAna)^86^. Specific thresholds were applied to different IF staining in the DiAna-segment module: a threshold of 5347 with a size larger than 1 pixel for CENPA, 2250 with a size larger than 1000 pixels for TEAD1 condensates, and 8405 with a size larger than 30 pixels for TRF2. Using the DiAna-analysis module, the minimal distance of each TEAD1 condensates to CENPA or TRF2 was determined. Distance histograms, visualizations, and statistical analyses were conducted using GraphPad Prism.

### Imaris 3D object surface classification and analysis

Nucleus, small TEAD1 condensate (STC), large TEAD1 condensate (LTC), YAP condensate (YAP), and *CYR61* intron-RNA FISH signal are identified by surface detection module using Imaris 10.2.0 (Bitplane) with absolute threshold of different channels manually set the same across images. STC and YAP surfaces were filtered by number of voxels 35-1000, LTC surfaces were filtered by number of voxels above 3000. Surface-to-surface distance analysis was performed by enabling shortest distance algorithm (Imaris). TEAD1 nuclear intensity was measured using nuclear 3D mask generated using Hoechst channel. *CYR61* RNA-FISH count and intensity were collected from averaged and specific values of build-in surface statistics.

### Live cell imaging, Drug Treatment, Fluorescence recovery after photobleaching (FRAP)

The UOK342 HaloTag-TEAD1 CRISPR knock-in cells were plated into 24-well glass bottom plate (Cellvis, P96-1.5H-N) for drug treatment and imaging the following day. Before drug treatment, the cells were labeled 10 nM JFX554 HaloTag ligand for 30 minutes. Then, the media was replaced with FluoroBrite DMEM Complete Medium (Gibco, A1896701) supplemented with 10% fetal bovine serum (FBS; Gibco, 26140079), 2 mM GlutaMAX-l (Gibco, 35050061), 1x NEAA (Gibco, 11140050). Live-cell images of individual cells were taken before and after treatments of 1% 1,6-hexandiol (Sigma-Aldrich, 88571) and DMSO (Corning, 01023017) for 30 minutes with 2 minutes interval.

UOK342 EGFP-TEAD1 stable cell line was plated into 8-well chambered cover glass (Cellvis, C8-1.5H-N) for FRAP experiment. Fluorescence bleaching was done on nuclear TEAD1 condensates using bleach mode in Zen software. The region of interest was selected on either the entire condensates or part of it using a rectangular box of approximately 1μm x 1μm in half-FRAP and 2μm x 2μm in full-FRAP. At least twenty iterations of bleaching were done with a 488nm Argon laser at 100% power. ∼140 rounds of imaging were performed before and after bleaching until fluorescence signal plateau, respectively, with an interval of 410ms. FRAP analysis was performed as previous studies. In short, for each time point, raw fluorescence intensities (*I*) from the bleached ROI, an unbleached ROI, a background ROI, and a reference region were measured. For full-bleach FRAP, bleached intensities (*FRAP_Full Bleach_*) were first computed by standard normalization *FRAP_Full Bleach_* = (*I_Background_ - I_Background_*)/(*I_Reference_ - I_Background_*) For half-bleach FRAP, bleach (*I_B_*) and non-bleached (*I_NB_*) intensities were further normalized to correct for an artifactual intensity drop in the non-bleached trace immediately after bleaching. Both traces were offset by a constant equal to the difference between the first two post-bleach nonbleach points (*I_I·B_* and *I_I·NB_*), yielding adjusted signals. Signals were then scaled by the relative ROI areas to account for differing region sizes, 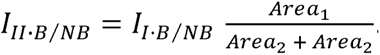. Next, traces were normalized to the pre-bleach and immediate post-bleach levels to obtain *I_III·B⁄NB_*. The immobile fraction was estimated from the terminal plateau by averaging the last three time points and comparing the non-bleached and bleached signals, and the recovery curve was corrected accordingly to generate the final normalized non-bleach trace and bleached trace (*I_IV·B⁄NB_*).The t_1/2_ was calculated using frapplot^87^, with fit formula for one diffusion component.

### Hyper/Hypo-osmotic stress treatment

UOK342 HaloTag-TEAD1 knock-in cells were plated into 8-well chambered cover glass plate 48 hours before treatments and replaced with FluoroBrite DMEM Complete Medium (Gibco, A1896701) with no FBS 1 hour before imaging. Cells were exposed to hyper-osmotic stress by adding 10x filter-sterilized 2M D-Sorbitol (Sigma, S1876), NaCl (RPI, 7647-14-5) dissolved in the FluoroBrite DMEM medium to a final concentration 0.2M or vehicle control. Hypo-osmotic stress was done by adding 50% v/v of filter-sterilized Molecular Biology Grade Water (Corning, 46-000-cv). Cells were imaged before and after treatment for 30 minutes with 2 minutes interval. To collect RNA from UOK342 TEAD1–HaloTag CRISPR knock-in cells, parental UOK cell lines and HK2 cells, cells were cultured similarly and collected 3 hours post 0.2M Sorbitol treatment or vehicle control.

### Western Blot

Cells were seeded on 6-well culture plates (FALCON, 353046), and plasmid transfection using Lipofectamine 3000 was done the day before cell lysis. After 20 hours of transfection, cells were washed ice-cold PBS, then add 100uL of lysis buffer (50mM Tris-HCl pH=8, 100nM NaCl, 1% IGEPAL, 0.5% Sodium deoxycholate) with 1X protease inhibitor (cOmpleteTM, Mini, EDTA-free Protease Inhibitor Cocktail). Cells were allowed to lyse on ice for 20 min and collect supernatant after centrifugation at 21,000 x g for 30 min at 4°C. Lysis was later added 1X loading buffer (10% glycerol, 2% SDS, 0.05M Tris-HCl pH=6.4, 0.01M 2-Mercaptoethanol, 0.6% bromophenol blue), and denatured at 90°C for 10 mins. 10uL mixture was later added to SurePAGETM 4-12% Bis-Tris gels (GenScript, #M00654). Protein was transferred to nitrocellulose membrane (LICOR, P/N 926-31090) by wet transfer, and blocked with 5% non-fat milk in TBS buffer. Membrane was incubated overnight at 4°C with 5% non-fat milk in TBS-T milk with primary antibodies (**Supplementary Table 2**). After washing three times in TBS-T, membranes were incubated at room temperatures for 1 h with secondary antibodies. Protein amounts were detected via LICOR Odyssey® M Imagers. Bands were quantified using FIJI.

### RT-qPCR and RNA-sequencing

Total RNA was isolated from HK2 and UOK cells lines using the Direct-zol RNA MiniPrep kit (Zymo Research, R2052) and converted to complementary DNA using the High-Capacity RNA-to-cDNA reverse transcription kit (Thermo Fisher, 4387406). The RT-qPCR was carried out on a QuantStudio 3 Real-Time PCR Instrument using PowerUp SYBR Green Master Mix (Thermo Fisher, A25742). Primers were purchased from Integrated DNA Technologies (IDT), and their sequences are detailed in **Supplementary Table 1**. mRNA levels of each gene were normalized to those of GAPDH.

RNA extraction for RNA-sequencing is performed similarly. RNA-seq was performed as a service at Novogene. In brief, mRNA samples were enriched using oligo(dT) beads and then used to prepare strand-specific cDNA libraries. Sequencing was done on Illumina Novaseq PE150. Individual reads are mapped to human genome hg38 by Hisat 2 v2.0.5. Featurecount in Rsubread was used to quantify gene expression level, and DESeq2Rpackage is used for differential expression analysis. Gene Ontology (GO) enrichment analysis of differentially expressed genes was used using DAVID^88^.

### ChIP-sequencing

Chromatin immunoprecipitation (Ch-IP) was performed based on previous studies^89^. In general, 10^8^ cells are used per each ChIP-seq experiment. HK2 and UOK cells were first treated with 1% formaldehyde in UOK media, followed by the addition of 0.125M glycine to halt cross-linking. The cells, washed in ice-cold PBS, were then lysed in cell lysis buffer, with subsequent nutation at 4°C. Cells were treated by Dounce homogenization on ice, followed by centrifugation at 500 x g for 10min. The supernatant was discarded, and the pellets were resuspended using nuclear lysis buffer. Nuclear lysates were then sheared by Covaris E220 to achieve an average chromatin length of 300-400 bp. The protein/chromatin complexes were captured using antibodies specific to the TEAD1, H3K9me3, YAP1, H3K27ac, and mouse IgG, and the mixtures were incubated overnight on the nutator at 4°C. The antibody/chromatin complexes were later captured by magnetic protein G beads and the washed and eluted antibody/chromatin complex using elusion buffer. After reversal of formaldehyde cross-linkes, DNA was purified with ChIP DNA Clean & Concentrator^TM^ (ZYMO research, D5205) and library preparation was performed using NEBNext^®^ Multiplex Oligos for Illumima^®^ (NEB, 10129566).

### ChIP-seq data analysis

ChIP-seq data were processed using the following pipeline. Raw sequencing reads in FASTQ format were quality-checked using FastQC. For identifying TEAD1/YAP co-occupy region, YAP and H3K27ac rReads were aligned to the human reference genome (hg38) using Bowtie2 (v2.3.5.3) with default parameter. The resulting BAM files were used to generate coverage tracks using *bamCoverage* from the *deepTools* suite^90^, with normalization of counts per million (CPM). Gene tracks are visualized using IGV 2.16.2.

Peaks for YAP and H3K27ac ChIP-seq were identified using MACS2 with default narrow peak settings and corresponding input controls (HK2 and UOK121 1% input) used to reduce background signal. A false discovery rate (FDR) cutoff of q < 0.01 was applied for peak selection. To identify putative co-regulatory TEAD1/YAP regions, overlapping peaks between TEAD1 and YAP datasets were identified using the *bedintersect* function in BEDTools. The resulting intersected peak regions were used for motif enrichment analysis via MEME-ChIP from the MEME Suite. Gene Ontology (GO) enrichment analysis of intersected peak regions was done using DAVID ^86^.

To investigate the genomic context of TEAD1 condensates, sequenced reads were mapped with Bowtie2 using repetitive mapping (-k 50). *SICER2* ^91^ was used for peak calling to identify broad enrichment regions characteristic of H3K9me3-marked heterochromatin. Peaks for TEAD1 and H3K9me3 ChIP-seq were identified using *SICER2* with broad peak parameter (-w 1000 rt 1 –fdr 0.05 -g 3000 –e 1000) and input controls with overlapped mapping (-k 50) ChIP-seq were selection. To identify TEAD1/H3K9me3 regions, overlapping peaks between the two were identified using the *bedintersect* function in BEDTools, and centromere regions are eliminated using *bedsubtract* function. To compare these co-bound regions across cell lines, we employed *computeMatrix*^92^ to quantify signal intensity at TEAD1 binding across all four cell lines. Heatmaps of TEAD1 signal were generated using *plotHeatmap* to visualize TEAD1 binding profiles in each cell line in TEAD1/H3K9me3 co-bound regions in condensates-positive cell lines (UOK121 and UOK342).

### Motif Discovery and Spacing Calculation

To quantify whether motif occurrences are enriched near centromeres, we analyzed genome-wide motif hit coordinates. BED files for MCAT (TEA transcription factor binding motif) and AP-1 (JUN/FOS transcription factors binding motif) were pre-build using FIMO^93^. Only primary hg38 chromosomes (chr1–chr22, chrX, chrY) were retained, and each motif hit was presented by its genomic midpoint. Centromere annotations collapsed per chromosome by taking the minimum start and maximum end. For each chromosome, we defined a pericentromeric interval as the centromere region expanded by ±4 Mb (bounded by chromosome ends). We then partitioned each chromosome into consecutive 1 Mb bins and counted motif midpoints per bin using a histogram-based approach; counts were converted to densities (hits per Mb) for each bin. Motif density distributions between pericentromeric and non-pericentromeric bins were compared using a one-tailed Welch’s t-test. To calculate the cumulative fraction (ECDF) within each chromosome, motif midpoints were de-duplicated to remove identical positions, sorted, and the distances between consecutive midpoints were computed (bp) using the first difference; each distance was assigned a genomic position at the midpoint of the corresponding gap.

### Binding and Expression Target Analysis (BETA)

To evaluate the regulatory function of TEAD1, we performed Binding and Expression Target Analysis using the BETA basic module^53^. ChIP-seq peak regions for TEAD1 were used as input along with RNA-seq differential expression results from UOK121 and UOK342 cells. Differentially expressed genes were identified based on log□ fold change thresholds, and categorized as upregulated, downregulated, or unchanged. The distance between TEAD1 peaks and target genes was defined using regions identified by the intersection of TEAD1 and H3K9me3 ChIP-seq peaks (via BEDtools), reflecting chromatin context-specific binding. BETA computed regulatory potential scores for each gene based on binding site proximity and density and assessed whether TEAD1 functions primarily as an activator or repressor using a Kolmogorov–Smirnov (KS) test to compare cumulative distributions of gene categories.

### TEAD1 expression, purification, and labelling

T7Express cells (NEB) were transformed with the pSAT vector encoding His-6, SUMO, TEAD1 (residues 1-426). Cells were grown in TB with 200 mg/mL ampicillin. Expression was induced with 0.5mM Isopropyl-β-D-1-thiogalactopyranoside (IPTG) at OD600=2.0, and cells were left shaking overnight at 20oC. The cell pellet was resuspended in 100mL of lysis buffer (50mM Tris pH7, 2M NaCl, 5mM b-ME, 5% glycerol, 10mM imidazole, supplemented with EDTA-free Protease inhibitor cocktail (Sigma) and 1mM phenylmethylsulfonyl fluoride (PMSF)), and lysed with a microfluidizer. Clarified lysates was incubated with Ni-charged Profinity IMAC resin (BioRad) for 30 minutes at 4oC. Resin was collected and then washed with 100mL of lysis buffer, followed by 100mL of lysis buffer supplemented with 15mM MgCl2, and 5mM ATP, and then 100mL of 50mM Tris pH 7, 500mM NaCl, 5mM b-ME, 5% glycerol, 20mM imidazole. Protein was eluted with 250mM imidazole and dialyzed overnight at 4oC into 50mM Tris7, 400mM NaCl, 5mM b-ME, 5% glycerol and the H6-SUMO tag removed by incubation with SENP (made in-house). Protein was pelleted with 40% NH4SO4, resuspended in 5ml of dialysis buffer and further purified with size exclusion chromatography (SEC) using a Superdex 75 16/600 column (GE) equilibrated in dialysis buffer. The protein was concentrated to 65uM and flash frozen (**Extended Data Fig. 7e**). Labelled TEAD1 was made as above except the dialysis buffer contained 10mM Hepes pH 7.4, was tagged with Alexa Fluor 488 TFP ester (Invitrogen) according to manufacturer’s instructions after gel-filtration, and then flash frozen. The final degree of labeling was determined to be 0.5 based on absorbance, as defined in the manual.

### Phase separation of recombinant TEAD1

DNA oligonucleotides were synthesized by IDT with the sequences listed in **Supplementary Table 3**. Equimolar amounts of complimentary oligonucleotides were resuspended at 1mM in 10mM Tris pH 8.0, 100mM NaCl and incubated at 95°C for 5 minutes in a heat block that was then cooled to room temperature overnight. Unlabeled and labeled TEAD1 were mixed for a final ratio of 10:1. 5µM of mixed TEAD1 was added to annealed dsDNA containing the indicated concentrations and sequences in 20mM Tris pH 7, 150mM NaCl, and 5mM MgCl2. Glass slides and coverslips were cleaned with 5% Hellmanex (Hellma Analytics), rinsed three times with water, then coated with 1% BSA solution for 1hour at room temperature, rinsed with water again, and left to dry. SecureSeal Imaging Spacers (Grace Bio-Labs) were placed on each coverslip. 3µl of TEAD1-DNA solution was added to each well. Wells were sealed and imaged using a Zeiss LSM 900 at 488nm. TEAD1 condensates were identified based on fluorescence intensity, size, and circularity using Fiji^84^. Six random locations were analyzed per condition, and each condition replicated three times. Data were plotted and one-way ANOVA test was performed using GraphPad Prism version 10.6.0 for Windows (GraphPad Software, Boston, Massachusetts USA).

## Proximity-based proteomics experiments

### IF confirmation of biotinylation of TEAD1 condensates components

Cells were plated on glass coverslips in a 24-well plate and transfected the next day at 70% confluency using lipofectamine 3000 Transfection reagent (L3000015; ThermoFisher), followed by 24-hour incubation. To image nuclear localization of the construct miniTurbo-TEAD1-HaloTag, cells were stained with JF646-HaloTag ligand (100nM; Janelia Materials) for 30 min before fixation. Cells were treated with 500µM Biotin in fresh growth media for 10 min at 37°C. Immediately after incubation, cell were washed with ice-cold DPBS for three times and fixed with 4% paraformaldehyde in PBS for 15 min on ice, permeabilized with ice-cold methanol at -20°C for 5 min, and blocked with 1% wt/v BSA in DPBS at 4°C for 30min. Primary antibodies against TEAD1(610923; BD Biosciences) was incubated overnight at 4°C. Cells were then incubated with secondary antibody Alexa Fluor 488 Goat anti-mouse IgG (H+L) (a11001, ThermoFisher). Biotinylated proteins were stained using streptavidin-Alexa Fluor 568 (S11226; ThermoFisher) for 1hr at room temperature. The cells were then incubated with Hoecht 33342 at RT for 5 min and mounted on glass slides with Vestashield and sealed with nail polish. Images were Airyscan-processed with the 1071 Zeiss Zen software (standard, 2D processing).

### Confirmation of protein biotinylation using Western Blot

Cells were plated on 10cm Petri dishes and transfected the next day at 70% confluency using lipofectamine 3000 Transfection reagent (L3000015; ThermoFisher), followed by 24-hour incubation. Cells were treated with 500µM Biotin in fresh growth medium for 10min at 37°C, washed with ice-cold DPBS for three times, and collected using cell scrapers and centrifuged at 300xg, 4°C for 5 min. Whole cell lysates were collected in 400µl RIPA buffer containing 50 mM Tris pH 8 (T60050-1000.0, Research Products International), 140 mM NaCl (S23025-3000.0, Research Products International), 1% Igepal® CA-630 (56741-50ML-F, Sigma Aldrich), 0.1% Sodium deoxycholate (30970-25G, Sigma Aldrich), 0.1% w/v sodium dodecyl sulfate (SDS, L3771, Sigma-Aldrich), and cOmplete™ Mini Protease Inhibitor Cocktail (11836170001, Sigma Aldrich) for 20 min on ice and clarified by centrifugation at 13,000 xg, 4°C for 10min. The protein concentrations were measured by BCA assay. 175µg of proteins from the lysates were enriched for biotinylation using Pierce Streptavidin Magnetic beads (88816; ThermoFisher) and incubated overnight at 4°C on a rotator. The bead slurry was washed twice with RIPA buffer, once with 1.0M KCL, once with 50mM TEAB buffer, and twice with RIPA buffer. About 10µg of proteins from Input, Post IP supernatant and pulldown samples were loaded on to the gel for immunoblotting. Biotinylated proteins were stained using Streptavitin-IRDye680 (926-68079; LICOR). Other antibodies used were anti-TEAD1 antibody (610923; DB Biosciences), and anti-β-actin antibodies (AC004; ABClonal).

### Sample preparation of proteomic analysis with miniTurbo

UOK342 cells stably expressing miniTurbo-TEAD1-HaloTag, miniTurbo-TEAD1 H3Mut-HaloTag, and miniTurbo-HaloTag-NLS were cultured overnight in 15cm Petri dishes and treated with 500µM Biotin for 10min at 37°C. Cells were collected and nuclear protein extract is collected using Nuclear Extract Kit (Active Motif, 40010). 600ug Nuclear Protein Lysate collected. Added 60ul Magnetic Bead (Washed twice with RIPA Buffer). Top the mixture with additional RIPA Buffer to 1mL. Incubate on a rotator overnight at 4°C. After overnight incubation streptavidin beads were collected on the side of the 1.5 mL tube using a magnet. transfer the lysate solution into new tubes. Beads were washed 2x with 1 mL of RIPA lysis buffer, 1x with 1 mL of 1.0 M KCl, 1x with 1 mL of 50 mM TEAB buffer and 2x with Tris-HCl pH7.5.

### TCA/Acetone Precipitation

Nuclear lysates were buffered to pH 8.0 with 100mM Triethylammonium bicarbonate (TEAB) in 1.5 ml centrifuge tubes, then reduced using 25 ul of 50mM dithiothreitol for 1 hr. at 60 C. Samples were chilled on ice then alkylated using 25 ul of 50mM iodoacetamide for 15 minutes in the dark. Samples were then diluted on ice in 8x volume of 10% trichloroacetic acid in acetone w/v stored at -20 C. Samples were vortexed and incubated for at least 4 hours at -20 C. Samples were then pelleted by centrifugation at 16,000g for 10 minutes at 0 C. The supernatant was carefully removed leaving the pelleted protein on the walls of the tube. 500ul of neat acetone stored at -20 C. was added to the pelleted protein and vortexed. This was then incubated for at least 10 minutes at -20 C. before centrifugation at 16,000g once again and the supernatant carefully removed. The remaining acetone was allowed to evaporate for 10 minutes before addition of trypsin (Pierce) in 100mM TEAB at a ratio of 1/50 enzyme to protein and allowed to digest overnight at 37C. Digested peptides were lyophilized and re-constituted in 500µl RIPA buffer and incubated with 50µl Pierce Streptavidin Magnetic beads overnight at 4°C on a rotator. The bead slurry was washed as described in WB confirmation followed by two times washes with ultrapure water. The washed beads were air-dried at 4°C for 6 hr and stored at -20°C.

Eluting peptides from streptavidin linked magnetic beads. Biotinylated peptides bound to streptavidin linked magnetic beads were separated from the final wash buffer by placing the Eppendorf tubes on a magnetic holder. After removing and setting aside the final wash buffer, the following steps were performed in the fume hood. Neat HFIP (200ul Millipore/Sigma Cat#105228) was added to each tube and put on a shaker at 1000 rpm for 10 minutes. The bead slurry was then placed on a magnetic holder to recover the HFIP eluted peptides from the magnetic beads. HFIP extraction was performed twice and both peptide elutions were combined in a new Eppendorf tube. The combined extractions were evaporated to dryness in a speedvac and resuspended in 0.1% TFA. Samples were buffer exchanged using an Oasis HLB (Waters, Milford MA) with two washes of 0.1% TFA and eluted with 0.1% TFA, 60% acetonitrile into new tubes and dried by vacuum centrifugation prior to mass spectrometry analysis.

### Mass Spectrometry analysis

Tryptic peptides were analyzed by reverse-phase chromatography tandem mass spectrometry on a Vantage NEO UPLC interfaced with an Orbitrap Exploris 480 mass spectrometer (Thermo Fisher Scientific). Peptides were separated using a 2%–90% acetonitrile in 0.1% formic acid gradient over 90 min at 300 nl/min using an in-house packed 75-um ID x 15 cm column packed with ReproSIL-Pur-120-C18-AQ (2.4 µm, 120 Å bulk phase, Dr. Maisch). Survey scans of precursor ions were acquired from 375-1500 m/z at a resolution setting of 120,000. Precursor ions were isolated in data dependent mode with a 3 second cycle time using a 1.5 dalton isolation width and a 45s dynamic exclusion. Peptides were fragmented at a normalized HCD collision energy of 30 with a resolution setting of 30,000.

Data analysis and candidate protein selection:

Mass Spec data for all 6 files were processed together using Proteome Discoverer v3.1.1.93 (ThermoFisher Scientific) using Sequest with Match Between Runs enabled against the UniProt Homo sapiens database. The TEAD constructs were inserted at the end of the Human FASTA file. Search criteria included trypsin as the enzyme with two allowed missed cleavages, a 10-ppm precursor mass tolerance and a 0.02 Da fragment mass tolerance. Modifications included fixed carbamidomethylation on C, oxidation on M, deamidation on N or Q and biotin on K as variable modifications. Peptide identifications were filtered at the 1% FDR level using the Percolator Node in Proteome Discoverer. Abundance ratio and Adjusted p-Values are generated and visualized by volcano plot showing differentially enriched proteins. String Analysis was generated based on the pulled candidate list. Protein-protein interaction networks and GO enrichment were rendered with settings at high confidence (0.700). Clusters composed of fewer than four proteins were removed. See **Supplementary Table 4** for proteomic source data. The mass spectrometry proteomics data have been deposited to the ProteomeXchange Consortium via the PRIDE [1] partner repository with the dataset identifier PXD072858 and 10.6019/PXD072858

### Single-particle tracking

Single-particle tracking (SPT) of TEAD1-Halo molecules was performed on a Nikon Ti2-Eclipse Motorized TIRF System with a x100/NA 1.49 oil-immersion objective, a Hamamatsu ORCA Fusion BT CMOS camera system, and a Perfect Focus System to combat axial focus drift. A Tokai Hit system was used to keep the humidified incubation chamber at 37 °C with 5% CO2. Nikon NIS-Elements was used to control the microscope hardware. UOK342 TEAD1-Halo cells were plated in 8-well chambered cover glass (Cellvis, C8-1.5H-N) 48 hours before imaging. On the day of imaging, cells were stained with 10 nM JFX554 Halo ligand for 30 minutes in FluoroBrite DMEM Complete Medium (Gibco, A1896701) supplemented with 10% fetal bovine serum (FBS; Gibco, 26140079), 2 mM GlutaMAX-l (Gibco, 35050061), 1x NEAA (Gibco, 11140050). Images used for SPT were obtained at 16-bit (no binning), 300 ms exposure and 25% 561 nm laser (LUN-F XL) for 2000 frames at an image size of 512x512 pixels (0.065 μm/pixel resolution). Prior to imaging, a snapshot of the same field of view was taken using epifluorescence (10 ms exposure, 25% 550 nm channel D-LEDI) for segmenting the nucleus and thresholding a mask of TEAD1 condensates during analysis. Four independent replicates were performed with 8 cells imaged per replicate. To correct for photobleaching of the fluorophore during subsequent data analysis, individual H2B molecules were also imaged as they are stably bound on chromatin. UOK342 cells were plated and transiently transfected as previously described with PB-EF1-Halo-H2B 16 hours prior to staining with JFX554 Halo ligand and imaging. At least 5 cells were imaged per replicate under the same parameters.

### SPT Analysis

The protocol found in Yoshida & Chong (2024)^94^ was used for SPT analysis to determine the dissociation kinetics and residence time of long-binding TEAD1 molecules in- and outside of TEAD1 condensates. The steps and parameters used are briefly described here. The nuclear region in each image stack was first segmented using the epifluorescence snapshot as a mask. We used a custom-written ImageJ macro using Trainable Weka Segmentation to create a binary mask of the nucleus from the snapshot which was then manually verified. The segmented image stacks were then converted to TIFF format and used in subsequent analysis. Localization and tracking of single molecules were performed using SLIMfast, a GUI implementation of the MTT algorithm found in Teves et al. (2016)^95^, with the following parameters: localization error = 10-6, deflation loops = 0, detection box = 9 pixels, lag time = 300 ms, max. expected diffusion coefficient (μm2/s) = 1, max. # competitors = 1. ImageJ macros and MATLAB code available in Chong et al. (2018)^96^ were used to threshold and create a binary mask of TEAD1 condensates from the same nuclear snapshot and subsequently categorize trajectories as being inside a condensate, defined as spending over 50% of lifetime within the masked area. A custom-written ImageJ macro was also used to rotate and place the condensate mask randomly within the nucleus so that it remained the same area as the condensate but did not overlap with it. Trajectories that were inside the original condensate or inside the randomly-placed mask in the nucleoplasm were then compiled by replicate and fit to a bi-exponential decay model to extract the measured dissociation rate constant (kobserved) and residence time (1/k). To correct for photobleaching, the same analysis was performed for Halo-H2B molecules in the whole nucleus (without categorization into a condensate), which are largely immobile. Because they were transiently transfected, we observed a proportion of H2B molecules that behaved unbound and were highly dynamic. As a result, we filtered out tracks that were less than 4 frames long and used the remaining trajectories to calculate k of the H2B molecules (kpb). We subtracted kpb from kobserved to get the true koff of TEAD1 molecules. Photobleaching correction was done per replicate, where each replicate was corrected by the kpb of Halo-H2B movies taken on the same day. The corrected residence time was calculated by 1/koff. The koff and residence times were averaged and GraphPad Prism was used to perform two-tailed paired t test between in-condensate and masked nucleoplasm trajectories.

## Notes

https://www.ncbi.nlm.nih.gov/geo/query/acc.cgi?acc=GSE298316

### Competing Interest Statement

The authors have declared no competing interest.

### Summary of Updates

This version of the manuscript has been revised to include the bioinformatics analysis of the clustered MCAT repeats on the pericentromeric heterochromatin, in vitro phase separation experiments of TEAD1 with repeated MCAT, direct testing of the depot model, and new proteomics control experiments, etc.

## Reference

1 Banani, S. F., Lee, H. O., Hyman, A. A. & Rosen, M. K. Biomolecular condensates: organizers of cellular biochemistry. Nat Rev Mol Cell Biol 18, 285–298 (2017). 10.1038/nrm.2017.7

2 Shin, Y. & Brangwynne, C. P. Liquid phase condensation in cell physiology and disease. Science 357 (2017). 10.1126/science.aaf4382

3 Mittag, T. & Pappu, R. V. A conceptual framework for understanding phase separation and addressing open questions and challenges. Molecular Cell 82, 2201–2214 (2022). 10.1016/j.molcel.2022.05.018

4 Lu, Y., et al. Phase separation of TAZ compartmentalizes the transcription machinery to promote gene expression. Nat Cell Biol 22, 453–464 (2020). 10.1038/s41556-020-0485-0

5 Demmerle, J., Hao, S. & Cai, D. Transcriptional condensates and phase separation: condensing information across scales and mechanisms. Nucleus 14, 2213551 (2023). 10.1080/19491034.2023.2213551

6 Hnisz, D., Shrinivas, K., Young, R. A., Chakraborty, A. K. & Sharp, P. A. A Phase Separation Model for Transcriptional Control. Cell 169, 13–23 (2017). 10.1016/j.cell.2017.02.007

7 Boija, A., et al. Transcription Factors Activate Genes through the Phase-Separation Capacity of Their Activation Domains. Cell 175, 1842-+ (2018). 10.1016/j.cell.2018.10.042

8 Muzzopappa, F., et al. Detecting and quantifying liquid-liquid phase separation in living cells by model-free calibrated half-bleaching. Nat Commun 13, 7787 (2022). 10.1038/s41467-022-35430-y

9 McSwiggen, D. T., et al. Evidence for DNA-mediated nuclear compartmentalization distinct from phase separation. Elife 8 (2019). 10.7554/eLife.47098

10 Cai, D., et al. Phase separation of YAP reorganizes genome topology for long-term YAP target gene expression. Nat Cell Biol 21, 1578–1589 (2019). 10.1038/s41556-019-0433-z

11 Sabari, B. R., et al. Coactivator condensation at super-enhancers links phase separation and gene control. Science 361 (2018). 10.1126/science.aar3958

12 Cho, W. K., et al. Mediator and RNA polymerase II clusters associate in transcription-dependent condensates. Science 361, 412–415 (2018). 10.1126/science.aar4199

13 Chong, S., et al. Tuning levels of low-complexity domain interactions to modulate endogenous oncogenic transcription. Mol Cell 82, 2084–2097 e2085 (2022). 10.1016/j.molcel.2022.04.007

14 Song, L., et al. Hotspot mutations in the structured ENL YEATS domain link aberrant transcriptional condensates and cancer. Mol Cell 82, 4080–4098 e4012 (2022). 10.1016/j.molcel.2022.09.034

15 Cai, D. F., Liu, Z. & Lippincott-Schwartz, J. Biomolecular Condensates and Their Links to Cancer Progression. Trends Biochem Sci 46, 535–549 (2021). 10.1016/j.tibs.2021.01.002

16 Spannl, S., Tereshchenko, M., Mastromarco, G. J., Ihn, S. J. & Lee, H. O. Biomolecular condensates in neurodegeneration and cancer. Traffic 20, 890–911 (2019). 10.1111/tra.12704

17 Linehan, W. M. The Genetic Basis of Kidney Cancer: Implications for Management and Use of Targeted Therapeutic Approaches. Eur Urol 61, 896–898 (2012). 10.1016/j.eururo.2012.02.022

18 Linehan, W. M. & Ricketts, C. J. The Cancer Genome Atlas of renal cell carcinoma: findings and clinical implications. Nat Rev Urol 16, 539–552 (2019). 10.1038/s41585-019-0211-5

19 Zheng, Y. & Pan, D. The Hippo Signaling Pathway in Development and Disease. Dev Cell 50, 264–282 (2019). 10.1016/j.devcel.2019.06.003

20 Fu, M., et al. The Hippo signalling pathway and its implications in human health and diseases. Signal Transduct Target Ther 7, 376 (2022). 10.1038/s41392-022-01191-9

21 Dong, J., et al. Elucidation of a universal size-control mechanism in Drosophila and mammals. Cell 130, 1120–1133 (2007). 10.1016/j.cell.2007.07.019

22 Zhao, B., et al. Inactivation of YAP oncoprotein by the Hippo pathway is involved in cell contact inhibition and tissue growth control. Gene Dev 21, 2747–2761 (2007). 10.1101/gad.1602907

23 Lei, Q. Y., et al. TAZ promotes cell proliferation and epithelial-mesenchymal transition and is inhibited by the hippo pathway. Mol Cell Biol 28, 2426–2436 (2008). 10.1128/Mcb.01874-07

24 Sourbier, C., et al. Targeting loss of the Hippo signaling pathway in NF2-deficient papillary kidney cancers. Oncotarget 9, 10723–10733 (2018). 10.18632/oncotarget.24112

25 Harvey, K. F., Zhang, X. & Thomas, D. M. The Hippo pathway and human cancer. Nat Rev Cancer 13, 246–257 (2013). 10.1038/nrc3458

26 Hao, S., et al. YAP condensates are highly organized hubs. iScience 27 (2024). 10.1016/j.isci.2024.109927

27 Wang, L., et al. Multiphase coalescence mediates Hippo pathway activation. Cell 185, 4376–4393 e4318 (2022). 10.1016/j.cell.2022.09.036

28 Bonello, T. T., et al. Phase separation of Hippo signalling complexes. EMBO J 42, e112863 (2023). 10.15252/embj.2022112863

29 Zhu, T., et al. Endogenous YAP/TAZ partitioning in TEAD condensates orchestrates the Hippo response. Mol Cell 85, 3425–3442 e3410 (2025). 10.1016/j.molcel.2025.08.014

30 Anglard, P., et al. Molecular and cellular characterization of human renal cell carcinoma cell lines. Cancer Res 52, 348–356 (1992).

31 Hao, S., et al. YAP condensates are highly organized hubs. iScience, 109927 (2024).

32 Boeynaems, S., et al. Protein Phase Separation: A New Phase in Cell Biology. Trends Cell Biol. 28, 420–435 (2018). 10.1016/j.tcb.2018.02.004

33 Roeder, R. G. & Rutter, W. J. Multiple forms of DNA-dependent RNA polymerase in eukaryotic organisms. Nature 224, 234–237 (1969). 10.1038/224234a0

34 Creyghton, M. P., et al. Histone H3K27ac separates active from poised enhancers and predicts developmental state. Proc Natl Acad Sci U S A 107, 21931–21936 (2010). 10.1073/pnas.1016071107

35 Shi, J. & Vakoc, C. R. The mechanisms behind the therapeutic activity of BET bromodomain inhibition. Mol Cell 54, 728–736 (2014). 10.1016/j.molcel.2014.05.016

36 Hnisz, D., et al. Super-enhancers in the control of cell identity and disease. Cell 155, 934–947 (2013). 10.1016/j.cell.2013.09.053

37 Zhang, H., Pasolli, H. A. & Fuchs, E. Yes-associated protein (YAP) transcriptional coactivator functions in balancing growth and differentiation in skin. Proceedings of the National Academy of Sciences 108, 2270–2275 (2011). doi:10.1073/pnas.1019603108

38 Quan, T., et al. Elevated YAP and its downstream targets CCN1 and CCN2 in basal cell carcinoma: impact on keratinocyte proliferation and stromal cell activation. Am J Pathol 184, 937–943 (2014). 10.1016/j.ajpath.2013.12.017

39 Kouzarides, T. Chromatin modifications and their function. Cell 128, 693–705 (2007). 10.1016/j.cell.2007.02.005

40 Eissenberg, J. C. & Elgin, S. C. HP1a: a structural chromosomal protein regulating transcription. Trends Genet 30, 103–110 (2014). 10.1016/j.tig.2014.01.002

41 Pirrotta, V. & Li, H. B. A view of nuclear Polycomb bodies. Curr Opin Genet Dev 22, 101–109 (2012). 10.1016/j.gde.2011.11.004

42 Matheson, L. & Elderkin, S. in Nuclear Architecture and Dynamics Vol. 2 (eds Christophe Lavelle & Jean-Marc Victor) 297–320 (Academic Press, 2018).

43 Verdaasdonk, J. S. & Bloom, K. Centromeres: unique chromatin structures that drive chromosome segregation. Nat Rev Mol Cell Bio 12, 320–332 (2011). 10.1038/nrm3107

44 Hall, L. E., Mitchell, S. E. & O’Neill, R. J. Pericentric and centromeric transcription: a perfect balance required. Chromosome Research 20, 535–546 (2012). 10.1007/s10577-012-9297-9

45 de Lange, T. Shelterin: the protein complex that shapes and safeguards human telomeres. Gene Dev 19, 2100–2110 (2005). 10.1101/gad.1346005

46 Anbanandam, A., et al. Insights into transcription enhancer factor 1 (TEF-1) activity from the solution structure of the TEA domain. Proc Natl Acad Sci U S A 103, 17225–17230 (2006). 10.1073/pnas.0607171103

47 Hwang, J. J., Chambon, P. & Davidson, I. Characterization of the transcription activation function and the DNA binding domain of transcriptional enhancer factor-1. EMBO J 12, 2337–2348 (1993). 10.1002/j.1460-2075.1993.tb05888.x

48 Li, Z., et al. Structural insights into the YAP and TEAD complex. Gene Dev 24, 235–240 (2010). 10.1101/gad.1865810

49 Lin, K. C., Park, H. W. & Guan, K.-L. Regulation of the Hippo Pathway Transcription Factor TEAD. Trends in Biochemical Sciences 42, 862–872 (2017). 10.1016/j.tibs.2017.09.003

50 Galli, G. G., et al. YAP Drives Growth by Controlling Transcriptional Pause Release from Dynamic Enhancers. Mol Cell 60, 328–337 (2015). 10.1016/j.molcel.2015.09.001

51 Zhu, C., et al. A Non-canonical Role of YAP/TEAD Is Required for Activation of Estrogen-Regulated Enhancers in Breast Cancer. Mol Cell 75, 791–806.e798 (2019). 10.1016/j.molcel.2019.06.010

52 Cebola, I., et al. TEAD and YAP regulate the enhancer network of human embryonic pancreatic progenitors. Nat Cell Biol 17, 615–626 (2015). 10.1038/ncb3160

53 Wang, S., et al. Target analysis by integration of transcriptome and ChIP-seq data with BETA. Nature Protocols 8, 2502–2515 (2013). 10.1038/nprot.2013.150

54 Farrance, I. K. G., Mar, J. H. & Ordahl, C. P. M-Cat Binding-Factor Is Related to the Sv40 Enhancer Binding-Factor, Tef-1. Journal of Biological Chemistry 267, 17234–17240 (1992).

55 Stein, C., et al. YAP1 Exerts Its Transcriptional Control via TEAD-Mediated Activation of Enhancers. PLoS Genet 11, e1005465 (2015). 10.1371/journal.pgen.1005465

56 Love, M. I., Huber, W. & Anders, S. Moderated estimation of fold change and dispersion for RNA-seq data with DESeq2. Genome Biol 15, 550 (2014). 10.1186/s13059-014-0550-8

57 Branon, T. C., et al. Efficient proximity labeling in living cells and organisms with TurboID. Nat Biotechnol 36, 880–887 (2018). 10.1038/nbt.4201

58 Cho, K. F., et al. Proximity labeling in mammalian cells with TurboID and split-TurboID. Nat Protoc 15, 3971–3999 (2020). 10.1038/s41596-020-0399-0

59 Alshareedah, I., Kaur, T. & Banerjee, P. R. Methods for characterizing the material properties of biomolecular condensates. Methods Enzymol 646, 143–183 (2021). 10.1016/bs.mie.2020.06.009

60 Alberti, S. The wisdom of crowds: regulating cell function through condensed states of living matter. Journal of Cell Science 130, 2789–2796 (2017). 10.1242/jcs.200295

61 Minshull, J., Blow, J. J. & Hunt, T. Translation of cyclin mRNA is necessary for extracts of activated Xenopus eggs to enter mitosis. Cell 56, 947–956 (1989). 10.1016/0092-8674(89)90628-4

62 Lockhead, S., et al. The Apparent Requirement for Protein Synthesis during G2 Phase Is due to Checkpoint Activation. Cell Rep 32, 107901 (2020). 10.1016/j.celrep.2020.107901

63 Ochneva, A., et al. Protein Misfolding and Aggregation in the Brain: Common Pathogenetic Pathways in Neurodegenerative and Mental Disorders. Int J Mol Sci 23 (2022). 10.3390/ijms232214498

64 Jonkers, J. & Berns, A. Oncogene addiction: Sometimes a temporary slavery. Cancer Cell 6, 535–538 (2004). 10.1016/j.ccr.2004.12.002

65 Serrano, M., Lin, A. W., McCurrach, M. E., Beach, D. & Lowe, S. W. Oncogenic ras provokes premature cell senescence associated with accumulation of p53 and p16INK4a. Cell 88, 593–602 (1997). 10.1016/s0092-8674(00)81902-9

66 Mooi, W. J. & Peeper, D. S. Oncogene-induced cell senescence--halting on the road to cancer. N Engl J Med 355, 1037–1046 (2006). 10.1056/NEJMra062285

67 Collado, M. & Serrano, M. Senescence in tumours: evidence from mice and humans. Nat Rev Cancer 10, 51–57 (2010). 10.1038/nrc2772

68 Sun, T. & Chi, J. T. Regulation of ferroptosis in cancer cells by YAP/TAZ and Hippo pathways: The therapeutic implications. Genes Dis 8, 241–249 (2021). 10.1016/j.gendis.2020.05.004

69 Magesh, S. & Cai, D. Roles of YAP/TAZ in ferroptosis. Trends Cell Biol 32, 729–732 (2022). 10.1016/j.tcb.2022.05.005

70 Charlier, C., Lamaye, F., Thelen, N. & Thiry, M. Ultrastructural detection of nucleic acids within heat shock-induced perichromatin granules of HeLa cells by cytochemical and immunocytological methods. J Struct Biol 166, 329–336 (2009). 10.1016/j.jsb.2009.03.002

71 Jolly, C., et al. In vivo binding of active heat shock transcription factor 1 to human chromosome 9 heterochromatin during stress. Journal of Cell Biology 156, 775–781 (2002). 10.1083/jcb.200109018

72 Biamonti, G. & Vourc’h, C. Nuclear stress bodies. Cold Spring Harb Perspect Biol 2, a000695 (2010). 10.1101/cshperspect.a000695

73 Raff, J. W., Kellum, R. & Alberts, B. The Drosophila Gaga Transcription Factor Is Associated with Specific Regions of Heterochromatin Throughout the Cell-Cycle. Embo Journal 13, 5977–5983 (1994). DOI 10.1002/j.1460-2075.1994.tb06943.x

74 Bhat, K. M., et al. The GAGA factor is required in the early Drosophila embryo not only for transcriptional regulation but also for nuclear division. Development 122, 1113–1124 (1996). 10.1242/dev.122.4.1113

75 Brown, K. E., et al. Association of transcriptionally silent genes with Ikaros complexes at centromeric heterochromatin. Cell 91, 845–854 (1997). Doi 10.1016/S0092-8674(00)80472-9

76 Klein, I. A., et al. Partitioning of cancer therapeutics in nuclear condensates. Science 368, 1386–1392 (2020). 10.1126/science.aaz4427

77 Silva, J. L., et al. Targeting Biomolecular Condensation and Protein Aggregation against Cancer. Chem Rev 123, 9094–9138 (2023). 10.1021/acs.chemrev.3c00131

78 Boija, A., Klein, I. A. & Young, R. A. Biomolecular Condensates and Cancer. Cancer Cell 39, 174–192 (2021). 10.1016/j.ccell.2020.12.003

79 Zhang, T., et al. FUS Regulates Activity of MicroRNA-Mediated Gene Silencing. Molecular Cell 69, 787–801.e788 (2018). 10.1016/j.molcel.2018.02.001

80 Deutsch, E. W., et al. The ProteomeXchange consortium at 10 years: 2023 update. Nucleic Acids Research 51, D1539–D1548 (2022). 10.1093/nar/gkac1040

81 Webster, M. R., et al. Wnt5A promotes an adaptive, senescent-like stress response, while continuing to drive invasion in melanoma cells. Pigment Cell & Melanoma Research 28, 184–195 (2015). 10.1111/pcmr.12330

82 Kraus, F., et al. Quantitative 3D structured illumination microscopy of nuclear structures. Nat Protoc 12, 1011–1028 (2017). 10.1038/nprot.2017.020

83 Sharifi-Sanjani, M., Meeker, A. K. & Mourkioti, F. Evaluation of telomere length in human cardiac tissues using cardiac quantitative FISH. Nat Protoc 12, 1855–1870 (2017). 10.1038/nprot.2017.082

84 Schindelin, J., et al. Fiji: an open-source platform for biological-image analysis. Nat Methods 9, 676–682 (2012). 10.1038/nmeth.2019

85 Bolte, S. & Cordelieres, F. P. A guided tour into subcellular colocalization analysis in light microscopy. J Microsc 224, 213–232 (2006). 10.1111/j.1365-2818.2006.01706.x

86 Heck, N., et al. A new automated 3D detection of synaptic contacts reveals the formation of cortico-striatal synapses upon cocaine treatment in vivo. Brain Struct Funct 220, 2953–2966 (2015). 10.1007/s00429-014-0837-2

87 Ding, G. (2019).

88 Sherman, B. T. et al. DAVID: a web server for functional enrichment analysis and functional annotation of gene lists (2021 update). Nucleic Acids Res 50, W216–W221 (2022). 10.1093/nar/gkac194

89 O’Geen, H., Frietze, S. & Farnham, P. J. Using ChIP-seq technology to identify targets of zinc finger transcription factors. Methods Mol Biol 649, 437–455 (2010). 10.1007/978-1-60761-753-2_27

90 Ramírez, F., et al. deepTools2: a next generation web server for deep-sequencing data analysis. Nucleic Acids Res 44, W160–W165 (2016). 10.1093/nar/gkw257

91 Zang, C., et al. A clustering approach for identification of enriched domains from histone modification ChIP-Seq data. Bioinformatics 25, 1952–1958 (2009). 10.1093/bioinformatics/btp340

92 Ramirez, F., Dundar, F., Diehl, S., Gruning, B. A. & Manke, T. deepTools: a flexible platform for exploring deep-sequencing data. Nucleic Acids Res 42, W187–191 (2014). 10.1093/nar/gku365

93 Grant, C. E., Bailey, T. L. & Noble, W. S. FIMO: scanning for occurrences of a given motif. Bioinformatics 27, 1017–1018 (2011). 10.1093/bioinformatics/btr064

94 Yoshida, S. R. & Chong, S. Single-Molecule Measurement of Protein Interaction Dynamics Within Biomolecular Condensates. JoVE, e66169 (2024). doi:10.3791/66169

95 Teves, S. S., et al. A dynamic mode of mitotic bookmarking by transcription factors. eLife 5, e22280 (2016). 10.7554/eLife.22280

96 Chong, S., et al. Imaging dynamic and selective low-complexity domain interactions that control gene transcription. Science 361, eaar2555 (2018). doi:10.1126/science.aar2555

